# Predicting perturbation effects from resting activity using functional causal flow

**DOI:** 10.1101/2020.11.23.394916

**Authors:** Amin Nejatbakhsh, Francesco Fumarola, Saleh Esteki, Taro Toyoizumi, Roozbeh Kiani, Luca Mazzucato

**Affiliations:** Center for Theoretical Neuroscience, Columbia University, New York, NY 10027, USA; Laboratory for Neural Computation and Adaptation, RIKEN Center for Brain Science, Wako, Saitama 351-0198, Japan; Center for Neural Science, New York University, New York, NY 10003, USA; Department of Psychology, New York University, New York, NY 10003, USA; Departments of Biology and Mathematics and Institute of Neuroscience, University of Oregon, Eugene, OR 97403, USA

## Abstract

A crucial challenge in targeted manipulation of neural activity is to identify perturbation sites whose stimulation exerts significant effects downstream (high efficacy), a procedure currently achieved by labor-intensive trial-and-error. Targeted perturbations will be greatly facilitated by understanding causal interactions within neural ensembles and predicting the efficacy of perturbation sites before intervention. Here, we address this issue by developing a computational framework to predict how single-site micorstimulation alters the ensemble spiking activity in an alert monkey’s prefrontal cortex. Our framework uses delay embedding techniques to infer the ensemble’s functional causal flow (FCF) based on the functional interactions inferred at rest. We validate FCF using ground truth data from models of cortical circuits, showing that FCF is robust to noise and can be inferred from brief recordings of even a small fraction of neurons in the circuit. A detailed comparison of FCF with several alternative methods, including Granger causality and transfer entropy, highlighted the advantages of FCF in predicting perturbation effects on empirical data. Our results provide the foundation for using targeted circuit manipulations to develop targeted interventions suitable for brain-machine interfaces and ameliorating cognitive dysfunctions in the human brain.

## I. INTRODUCTION

Cognition is an emergent property of collective interactions of neurons in large networks of cortical and subcortical circuits. Targeted manipulation of the brain to alter behavior will be greatly facilitated by understanding the causal interactions within these circuits. Successful examples of such manipulations include changing perceived motion direction by altering responses of direction selective neurons in area MT [1, 2], biasing object classification towards faces by altering responses of face selective neurons in the inferior temporal cortex [3–5], changing the value of a stimulus by altering neural responses in the anterior caudate [6], and controlling movements and postures by altering the activity of motor and premotor cortical neurons [7].

Successful behavioral manipulations depend on the identification of suitable perturbation sites, satisfying at least two requirements. The first is selectivity: the local neural population around the perturbation site should exhibit response properties bearing on the desired behavioral effect, e.g., motion direction selectivity in area MT [1, 2], face selectivity in face patches of inferotemporal cortex [3–5], or the locus of seizure in epilepsy [8]. The second requirement is efficacy: stimulation of the local population should exert some significant effect on the activity of the rest of the brain, and consequently on behavior. While selectivity of sensory and motor neurons may be estimated by recording neural activity in simple and well-defined tasks, selectivity tends to be quite complex or variable across tasks in many regions of the association cortex. Further, discovering efficacy is currently achieved by trial-and-error: many perturbations are performed until a site whose stimulation leads to a significant change in activity is located. As a result, current methods for targeted perturbations are labor intensive, time consuming, and often unable to generalize beyond the limited task set they are optimized for.

A promising alternative for predicting the efficacy of a potential perturbation site is to examine its functional connectivity within a local neural circuit. Intuitively, one expects that perturbing a node with strong functional connectivity to other nodes within a circuit may exert stronger effects than perturbing the nodes that are functionally isolated. Estimating the functional connectivity in cortical circuits is a central open problem in neuroscience [9], and currently-used methods, which are adapted from frameworks developed for generic multidimensional time series, are challenged by the specific properties of neural activity in the cortex. Cortical circuits comprise highly recurrent neural networks [10–13], where the notion of directed functional couplings is not obvious. Correlation-based methods [14] lack sufficient power when correlations are weak, as in most cortical circuits [15]. Granger causality, a widely used method [16–19], relies on the assumption of linear dynamics and thus it is challenged when the circuit’s dynamical properties are not well known. Transfer entropy [20], applicable to non-linear systems, requires large datasets hard to acquire in conventional experiments. Critically, both Granger- and entropy-based methods require stochasticity and are challenged in the presence of self-predictability in deterministic dynamics such as nonlinear couplings between variables [21]. Moreover, commonly encountered confounding effects such as phase delay [22] or common inputs [21] render these methods unreliable. It is thus of paramount importance to develop new theoretical tools. These new tools should be able to estimate causal functional interactions in the presence of common inputs and unknown non-linear dynamics. Further, to be useful for translational applications, they should accommodate extremely sparse sampling (i.e., from a small fraction of neurons in a circuit), typical of electrophysiological recordings in humans and monkeys.

We propose a novel approach to infer causal functional interactions using delay embedding methods, in particular, convergent cross-mapping (CCM) [21], a technique expressly developed to work in the sparse data regime [23, 24] and in the presence of noise, common inputs, and nonlinear couplings between variables [21] — hallmarks of cortical dynamics that explicitly violate the assumptions underlying many alternative methods (e.g., Granger). The key idea of CCM is that single observation variables in a complex dynamical system can be used to construct shadow manifolds with one to one correspondence with the attractor manifolds of the full system. While the powerful CCM framework, rigorously articulated in [25], has been successfully applied in ecology [21], and *in vitro* [21, 26] and ECoG neural activity [27], it has never been adapted to spiking activity *in vivo*. We build on CCM to develop new statistics, termed “Functional Causal Flow” (FCF), which captures the reconstruction accuracy of the response dynamics of units in an ensemble from the activity of the other units (Fig 1).

**FIG. 1.**
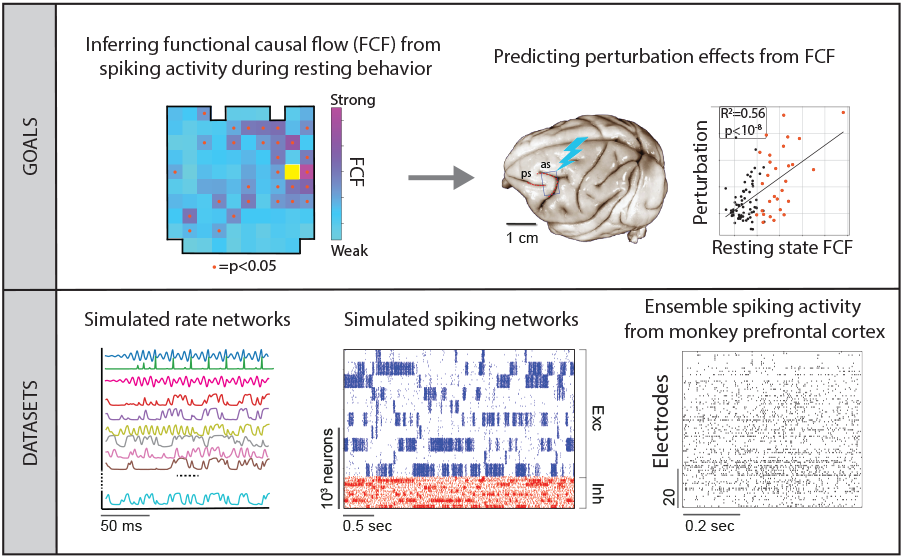
Conceptual summary. Top left: Functional causal flow (FCF) map inferred from the prefrontal cortex activity of a resting alert monkey. Each square represents an electrode of a 96-electrode Utah array (yellow square: microstimulated electrode; orange dots: electrodes with significant FCF that are functionally downstream to the stimulated one). Top right: Schematics of an electrical microstimulation experiment, where we use FCF to predict the magnitude of neural activity perturbations. The scatter plot shows the correlations of resting state FCF vs. perturbation effects on targets with significant (orange dots) and non-significant FCF (black dots). Bottom: We validated our method for predicting perturbation effects from resting state FCF in three different datasets: a chaotic rate network, a spiking network with cell-type specific connectivity, and a prefrontal cortical circuit in alert monkeys.

We characterize the effects of perturbations by introducing the concept of interventional connectivity, an observable that is agnostic to the underlying structural connectivity and only depends on responses to perturbations; and show that one can efficiently predict interventional connectivity solely based on FCF inferred from short spike trains. We perform a series of simulated experiments to determine applicability regimes of FCF. In these simulated experiments, we present strong challenges, such as noise and common inputs, yet retain features of biological plausibility from known cortical circuit dynamics. These simulations validate accuracy and efficiency of FCF. We then demonstrate that our method infers the causal flow of ensembles of neurons from sparse recordings of spiking activity, obtained from chronically implanted prefrontal multi-electrode arrays in awake, resting monkeys. Using the causal flow inferred during resting activity, we successfully predict the effect of electrical microstimulations of single sites on the rest of the circuit. Finally, we perform a detailed comparison between FCF and alternative methods used to infer functional interactions, including Granger causality, non-linear extensions of Granger causality, and transfer entropy. This critical comparison demonstrates the superior performance of FCF for both the simulated and empirical data. We have created and shared an open source tutorial software package for efficient estimation of FCF and alternative methods on simulated and empirical data, which can be easily and quickly run on conventional computers. In summary, our results highlight the advantages of deploying causal flow to guide perturbation experiments compared to traditional methods, opening the way for much more efficient protocols for targeted manipulations of cortical ensembles in primates and humans.

## II. RESULTS

### A. Uncovering the functional causal flow with delay embedding

To illustrate the concept and methods of functional causal flow (FCF) and establish its regime of validity, we performed a series of simulated experiments using ground truth data from recurrent network simulations. We chose two sets of neural networks (Fig. 1). In the first experiment, we simulated a continuous rate network comprising both feedforward and recurrent features in its structural connectivity, where we arbitrarily varied the noise levels and features to assess FCF robustness against different signal-to-noise ratios and common inputs. In this simulated experiment, we benchmarked FCF estimation against alternative causality indices such as Granger causality, its nonlinear extensions, and transfer entropy. In the second experiment, we simulated a cortical circuit model based on a spiking network with cell-type specific connectivity endowed with functional assemblies. This class of spiking models captures the intrinsic neural variability observed in various cortical areas across different tasks and behavioral states[28–32]. Calibrating our FCF inference on this spiking model served as a guide for our experimental data from alert monkeys.

We first examined a deterministic network Z, comprising N units arranged in two subnetworks X and Y, each endowed with their own local recurrent connectivity. Crucially, there are directed projections from X to Y with coupling strength *g*, but no feedback couplings from Y to X. Units in X represent a chaotic Rossler attractor and have strong all-to-all recurrent couplings, while units in Y have only sparse and weak recurrent couplings. We aimed to capture the intuitive idea that the upstream subnetwork X drives the activity of the downstream subnetwork Y (Fig. 2). It is well known from the theory of deterministic dynamical systems that one can (at least partially) reconstruct the N-dimensional attractor topology of a network of coupled units, represented by the vector time series of the activity of all units 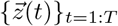, by using only the information encoded in the temporal trajectory of a single unit*{z*_*i*_(*t*)*}*_*t*=1:*T*_. From the mapping between the activity of the full network and the activity of a single unit, one can derive a map between the activity of the units themselves and (at least partially) reconstruct the activity of one unit*{z*_*i*_(*t*)*}*_*t*=1:*T*_ from the activity of a different unit*{z*_*j*_(*t*)*}*_*t*=1:*T*_, for *i* ≠ *j*. The reconstruction is possible whenever the two units are functionally coupled. This general property of dynamical systems is known as “delay embedding” [23, 24] and relies on a representation of network dynamics using “delay coordinates” (see Fig. 2A for details). This reconstruction has also been shown to be robust to noise in driven dynamical systems [33].

**FIG. 2.**
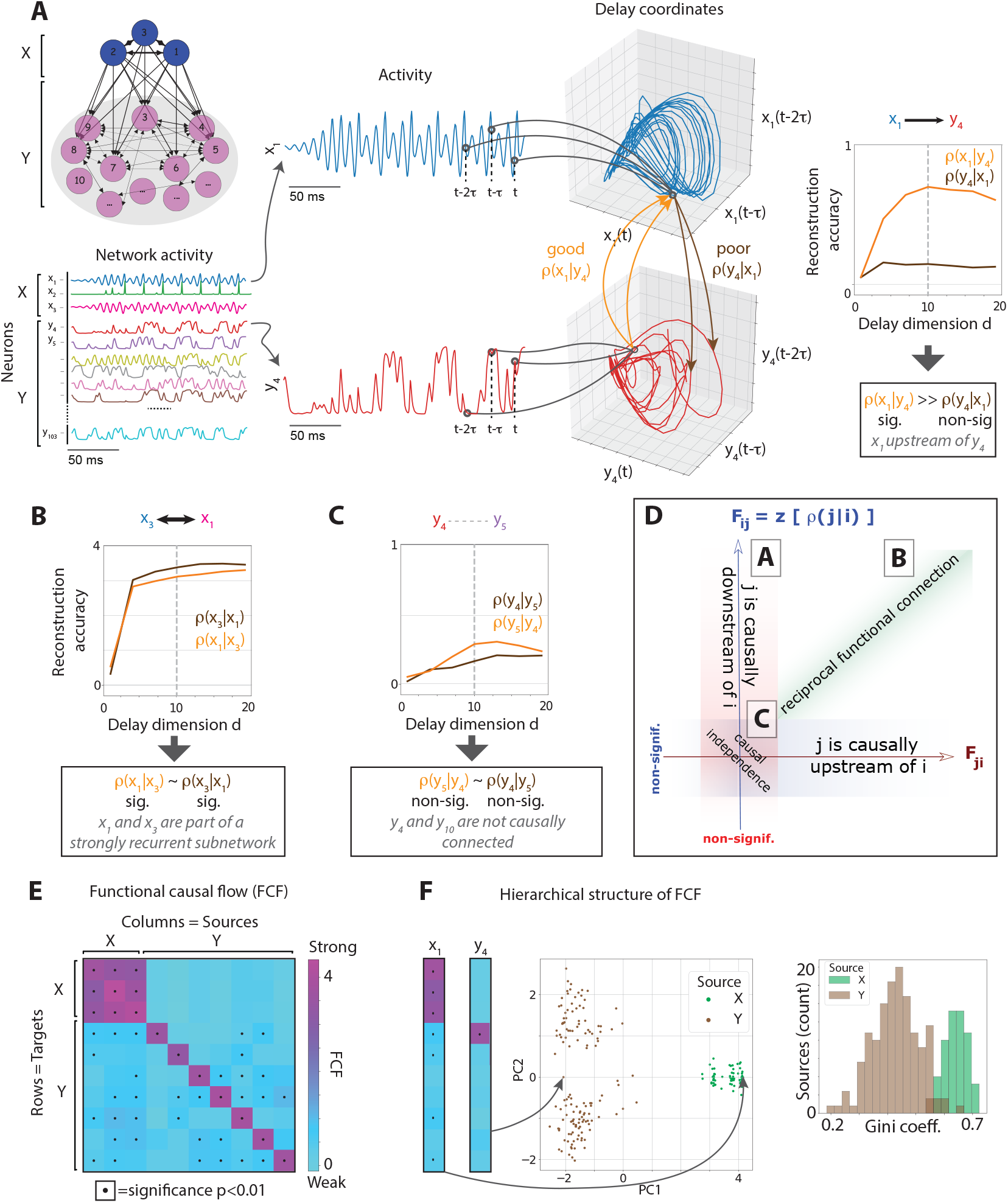
Functional causal flow. A) Left: Schematic of network architecture Z: two subnetworks X (blue nodes) and Y (pink nodes) with strong and weak recurrent couplings within the subnetworks, respectively, are connected via feedforward couplings from X to Y. Thickness of black arrows represents the strength of directed structural couplings. Center: Activity of units *y*_4_(*t*) (red, bottom) and *x*_1_(*t*) (blue, top) are mapped to the delay coordinate space *X*_1_ = [*x*_1_(*t*), *x*_1_(*t−τ*), …, *x*_1_(*t−* (*d−*1)*τ*)] and *Y*_4_ (right, *τ* = 4ms, see Fig S1A). Reconstruction accuracy increases with delay vector dimension *d* before plateauing. The reconstruction accuracy *ρ*(*x*_1_|*y*_4_) of upstream unit *x*_1_ given the downstream unit *y*_4_ is significant and larger than the reconstruction accuracy *ρ*(*y*_4_|*x*_1_) of *y*_4_ given *x*_1_ (non-significant). The FCF value *F*_41_ reveals a strong and significant causal flow from upstream node *x*_1_ to downstream node *y*_4_. B) The significant FCF between two units *x*_1_ and *x*_3_ within the strongly coupled subnetwork X reveal strong and significant causal flow between them, but no preferred directionality of causal flow. C) The non-significant FCF between two units *y*_4_ and *y*_5_ in the weakly coupled subnetwork Y suggests the absence of a causal relationship, matching our network design. D) Summary of the FCF cases in panels A, B, C (see table I). E) The FCF between 10 representative units sparsely sampled from the network (columns and rows represent source and target units, respectively; columns are sorted from functionally upstream to downstream units). F) The functional hierarchy in the network structure is encoded in the causal vectors. Left: PCA of columns of the FCF matrix. Each dot represents the causal vector of one source unit in a trial. Twenty trials with different initial conditions were simulated. The same X (green) and Y units (brown) of panel E are depicted. The separation of X and Y units is strongly preserved regardless of the initial condition. Right: Distribution of Gini coefficient of causal vectors.

**TABLE I.** Definitions and notations for the functional causal flow (FCF).

|  | FCF | Feature |
| --- | --- | --- |
| Upstream | $F_{ij} > 0$ & sig; $F_{ji}$ non-sig | $j$ is causally upstream of $i$. |
| Downstream | $F_{ij}$ non-sig; $F_{ji} > 0$ & sig | $j$ is causally downstream from $i$. |
| Reciprocal | $F_{ij} \sim F_{ji}$ & both sig | $i$ and $j$ are reciprocally functionally connected. |
| Independent | $F_{ij}, F_{ji}$ both non-sig | $i$ and $j$ are causally independent. |

We used convergent cross-mapping based on delay embedding to infer the FCF between all pairs of network units. We first considered the FCF between a unit *y*_*i*_ in the downstream subnetwork Y and a unit *x*_*j*_ in the upstream subnetwork X. The activity of unit *x*_*j*_ only depends on the other units in X, to which it is recurrently connected, but not on the units in Y, as there are no feedback couplings from Y to X. On the other hand, the activity of unit *y*_*i*_ depends both on the units in X, from which it receives direct projections, and on the other units in Y to which it is recurrently connected. In other words, *y*_*i*_(*t*) is causally influenced by units in both Y and X, whereas *x*_*j*_(*t*) depends only on other units in X. Thus, we expect that the reconstruction of *x*_*j*_(*t*) from *y*_*i*_(*t*) will be more accurate than the reconstruction of *y*_*i*_(*t*) from *x*_*j*_(*t*), because in the latter case the causal influence on *y*_*i*_ from the other recurrently connected units in *Y* is being neglected.

We tested our prediction by reconstructing the temporal series of unit *x*_*j*_(*t*) given *y*_*i*_(*t*) from the corresponding delay vectors [*x*(*t*), *x*(*t−τ*), …, *x*(*t−dτ* + *τ*)] and [*y*(*t*), *y*(*t−τ*), …, *y*(*t−dτ* + *τ*)] of dimension *d* with a step *τ*. Reconstruction accuracy was quantified as the Fisher z-transform *z*[*ρ*(*y*_*i*_|*x*_*j*_)] of the Pearson correlation *ρ*(*x*_*j*_|*y*_*i*_) between the *empirical* activity of the delay vector of unit *x*_*j*_ and its *predicted* activity obtained from the delay vector of unit *y*_*i*_. Whereas the Pearson correlation is bounded between *−*1 and 1, its Fisher z-transform is approximately normally distributed thus facilitating statistical comparisons [34]. The process was cross-validated to avoid overfitting (see Methods for details). Similarly, we estimated cross-validated reconstruction accuracy *z*[*ρ*(*y*_*i*_ *x*_*j*_)] of the temporal series of unit *y*_*i*_(*t*) given *x*_*j*_(*t*). As expected, reconstruction accuracy increased as a function of the dimensionality of the delay coordinate vector (i.e., how many time steps back we utilize for the reconstruction, Fig. 2A). The accuracy plateaued beyond a certain dimensionality (related to the complexity of the time series [26]), whose value we fixed for our subsequent analyses. Model selection for hyperparameters *d* (delay dimension) and *τ* (time step) depended on the specific datasets. For the continuous rate network considered here model selection yielded optimal values of *d* = 7 and *τ* = 4 ms (see Fig. S1A). For delay dimensions *d ≥*5, the sample size dependence of FCF plateaued at about 1500 ms (Fig. S1B).

We define the functional causal flow (FCF) from *x*_*j*_ to *y*_*i*_ as *F*_*ij*_ = *z*[*ρ*(*x*_*j*_|*y*_*i*_)] (see Methods and table I for a summary of our conventions). Columns of the FCF represent the *source* units, whose activity is being reconstructed, and rows represent the *target* units, whose activity is used for the reconstruction. We established statistical significance by comparing the FCF estimated from the empirical data with that estimated from surrogate datasets designed to preserve the temporal statistics of the network activity while destroying its causal structure [35] (see Fig. S2 and Methods for details). Surrogates were produced in three steps: first, nearest neighbors of a state were identified in the delay-embedding space; second, “twin” states were constructed for each state including all neighbors within a small distance; finally, surrogate trajectories were generated by temporally concatenating states from the same coarse-grained sets of twins, allowing for jumps backward or forward in time while preserving all large-scale nonlinear properties of the system.

Unlike the Pearson correlation *r*_*ij*_, which is a symmetric quantity, the FCF is a directed measure of causality. By comparing the value and significance of *F*_*ij*_ with *F*_*ji*_, we can establish the *directionality* of the functional relationship between *y*_*i*_ and *x*_*j*_, uncovering qualitatively different causal flow motifs (table I), as we explain in the next section.

### B. Functional causal flow uncovers hierarchical structures

In the example above, the reconstruction accuracy of *x*_*j*_ given *y*_*i*_ was significant and large, while that of *y*_*i*_ given *x*_*j*_ was not significant. In other words, while one can significantly reconstruct *x*_*j*_ with high accuracy from *y*_*i*_, because the latter receives information from the former, the opposite is not possible, matching predictions based on the simulated network architecture. We refer to *x*_*j*_ as being *causally upstream* to *y*_*i*_ in the functional causal flow of the network.

The notion of being causally upstream or downstream is an entirely *functional* relation and *a priori* different from the underlying anatomical coupling between units. We illustrate here two more examples from the network in Fig. 2 to reveal the variety of the relationships encoded in the FCF. First, we considered the FCF between *x*_1_(*t*) and *x*_3_(*t*) within subnetwork X (Fig. 2B). Because subnetwork X does not receive inputs from Y, it is causally isolated (i.e., its activity is conditionally independent from Y). Hence one can reconstruct the activity of one *x*_*i*_ unit from another with high accuracy, yielding large and significant FCF values. Fig. 2B shows large FCF both *x*_1_|*x*_3_ and *x*_3_|*x*_1_. This is a classic demonstration of the embedding theorem [23], ensuring accurate bidirectional reconstruction of variables mapping a chaotic attractor. The large and significant 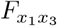 and 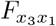 reveal that the unit pair has a strong reciprocal functional coupling, and the two units lie at the same level of the functional hierarchy. This is unlike the case of pairs *x*_*j*_, *y*_*i*_ described above, where a significant *F*_*ij*_ but a non-significant *F*_*ji*_ showed a strong directional coupling and a functional hierarchy [36]. As another qualitatively different pair, we considered two units *y*_4_ and *y*_5_ within subnetwork *Y*, whose units are only sparsely recurrently coupled. FCFs were not significant for this pair (Fig. 2C), suggesting that the two units are functionally independent, namely, their activities do not influence each other significantly. The taxonomy of causal flows are summarized in table I and Fig. 2D.

The variety of FCF features discussed so far suggests that, even though the FCF is a measure of pairwise causal interactions, it may reveal a network’s global causal structure. We thus analyzed the *N*-dimensional *causal vectors* 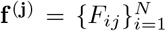, representing the reconstruction accuracy of unit *j* given the activity of each one of the targets *i*. The causal vector **f** ^(**j**)^ encodes the FCF from unit *j* to the rest of the network. A Principal Component analysis of the causal vectors from a sparse subsample of the network units (10 out of 103) revealed a clear hierarchical structure present in the network dynamics showing two separate clusters corresponding to the subnetworks X and Y (Fig. 2E-F). Thus, causal vectors revealed the global network functional hierarchy from sparse recordings of the activity.

We further quantified the hierarchical functional structure of causal vectors, measured by their Gini coefficients (Fig. 2F), which estimates the degree of inequality in a distribution. For example, a delta function, where all samples have the same value, has zero Gini coefficient, while an exponential distribution has a Gini coefficient equal to 0.5. In the absence of hierarchical structures, one would expect all targets from a given source unit to have comparable FCF values, yielding a low Gini coefficient. Alternatively, heterogeneity of FCFs across targets for a given source would suggest a network hierarchy with a gradient of functional connectivities, yielding a large Gini coefficient. For our simulated network, we found a large heterogeneity in the Gini coefficients of the causal vectors, capturing the functional hierarchy in the network. For comparison, when restricting the causal vectors to sources in either X or Y (green and brown bars in Fig. 2F, respectively), we found a clear separation with larger Gini coefficients for X sources and lower Gini coefficients for Y sources. This result shows that the feedforward structural couplings from X to Y introduce a hierarchy in the full network Z, encoded in the network causal vectors.

### C. Robustness of functional causal flow estimation to noise and common inputs

Neural circuits in vivo include multiple sources of variability including both private (e.g., Poisson variability in spike times) and shared variability (e.g., low-rank cofluctuations across the neural ensemble [37, 38]), where the latter may correlate to the animal’s internal state such as attention or arousal [39–42]. Shared sources of external variability represent common inputs, which notoriously pose strong challenges to existing methods to infer functional connectivity. Based on previous theoretical work, we expected our delay embedding framework to be reasonably resilient against these effects [33].

To quantify the robustness of inferred FCFs, we tested their changes as a function of the strength of a noisy input to subnetwork Y. When driving the network with either private noise (i.i.d. for each neuron, Fig. 3A) or common input (shared noise, namely, the same noise realizations across all neurons, Fig. 3B), we found that FCF inference degraded only when the signal-to-noise ratio (SNR) dropped below 0 dB (Fig. 3C, SNR is measured in logarithmic scale). However, for a wide range of SNRs, the FCF inference maintained its accuracy. The degradation caused by private noise did not remove the informativeness of FCFs for the tested range of SNRs down to −10 dB. Common external input in the form of shared noise became irrecoverably detrimental only for SNRs below −5. We thus conclude that causal flow estimates are robust to both noise and external time-varying inputs, two common sources of neural variability.

**FIG. 3.**
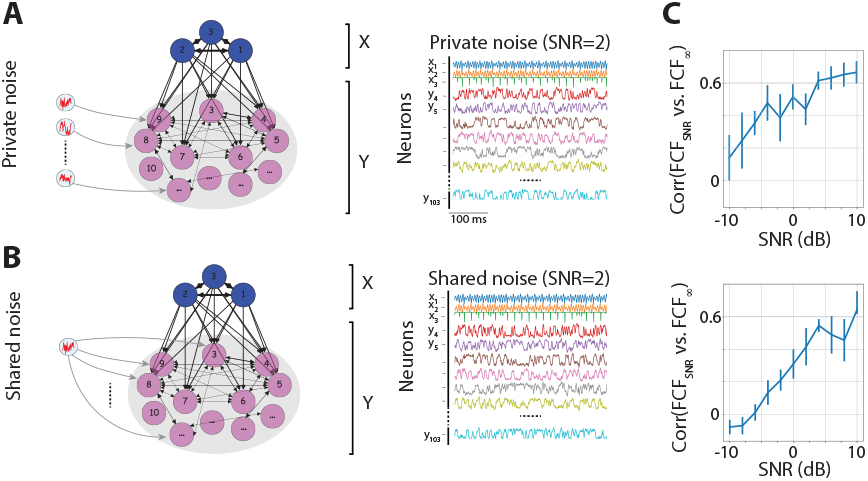
Robustness of inferred FCFs to time-dependent inputs. FCF is robust to both private noise (panel A, i.i.d. realizations of noise in each unit) or shared noise (panel B, a scalar noise source modulates all neurons). Same network as in Fig. 2. C) Correlation of FCF inferred from the noiseless simulations (FCF_*∞*_) with the FCF inferred from simulations with varying SNR (SNR, defined in units of dB as 10 log_10_[*σ*(*signal*)*/σ*(*noise*)], where *σ* =standard deviation). Top: private noise; bottom: shared noise.

We also explored another source of variability common in the experimental data: recordings across different sessions, which we modeled as changes in the initial conditions for the network dynamics. We performed multiple simulations of the same network where each simulation varied in regard to initial conditions, and thereby the sequence of recorded neural activity. We found that FCF was largely indistinguishable across the simulated sessions with different initial conditions.

Our validation results thus demonstrate that the datadriven discovery of functional causal flow is robust to noise. Another source of variability is the presence of unobserved units in the network, which we will address below (see Section “Causal flow from sparse recordings in spiking circuits”).

### D. Inferred causal flow predicts the effects of perturbation

Can we predict the effects of perturbations on network activity based on the FCF inferred from the unperturbed system? We hypothesized that the effects of stimulating a specific node on the other nodes of a network can be predicted by the causal flow inferred during the resting periods.

We simulated a perturbation protocol where we artificially imposed an external input on one source unit for a brief duration, mimicking electrical or optical stimulation protocols of cortical circuits (Fig. 4A). We estimated the stimulation effect on each target unit, by comparing the distribution of activity in each target in intervals preceding the stimulation onset and following its offset (Fig. 4B). We found that stimulation exerted complex spatiotemporal patterns of response across the target units, which we captured with a *perturbation vector*: 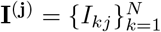, where *I*_*kj*_ is “interventional connectivity” between target unit *k* and source unit *j* (Fig. 4B). Interventional connectivity is quantified as the Kolmogorov-Smirnov statistics between pre- and post-stimulation spiking activity distributions over stimulated trials. In our simulations, stimulation effects across targets *k* strongly depended on the source unit *j* that was stimulated. Perturbation effects increased with stimulation strength for source-target pairs in *X→X* and *X→Y* and *Y→Y*, but not for pairs *Y→X*, consistent with the underlying structural connectivity lacking feedback couplings from *Y* to *X* (Fig. 4C). Can one predict the complex spatiotemporal effects of stimulation solely based on the FCF inferred during resting activity?

**FIG. 4.**
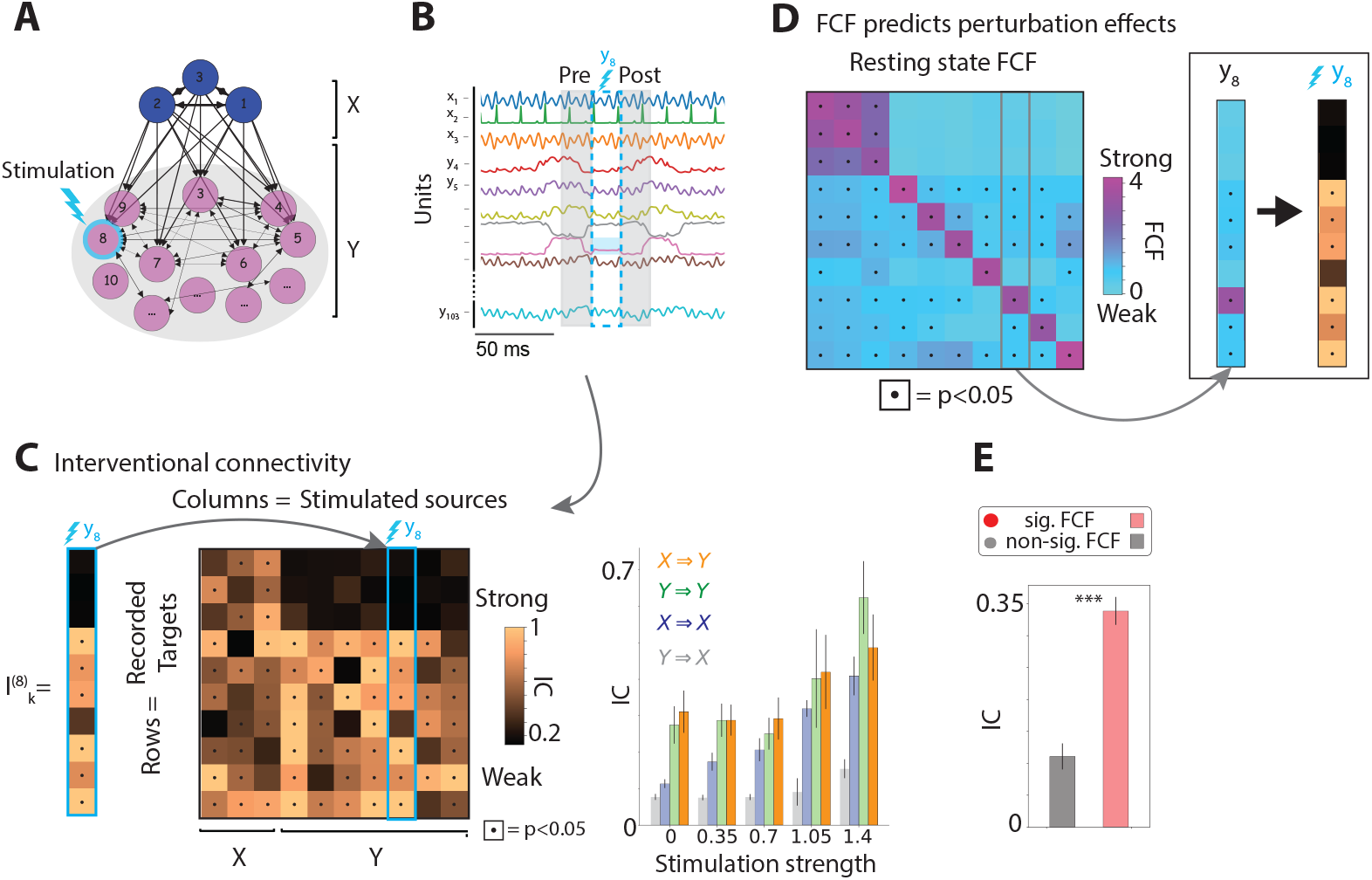
Functional causal flow predicts perturbation effects. A) Perturbation protocol: single nodes (e.g., unit *y*_8_) are stimulated with a pulse of strength *S* lasting for 100ms. B) Perturbation effects on target units are estimated by comparing the activity immediately preceding the onset and following the offset of stimulation to calculate Interventional Connectivity (IC). C) IC is the Kolmogorov-Smirnov test statistics for the distribution of preand post-stimulation activity of the target units. Left: IC matrix for 10 representative units. Black dots represent significant effects (*p <* 0.05). The effects of stimulating one source *i* on all targets *k* is encoded in the perturbation vector **I**^(**i**)^. Right: Perturbation effects increase with the stimulation strength *S* for source-target pairs of *X→X, Y→Y*, and *X→Y*, but not *Y→X*, reflecting the absence of feedback structural couplings from *Y* to *X*. Error bars are s.e.m.. D) For each source, its causal vector (e.g., column of the resting state FCF for source unit *y*_8_) is compared with the perturbation vector (columns of the interventional connectivity matrix), revealing that FCF predicts perturbation magnitudes. E) target units with significant resting state FCF had a larger response to stimulation, compared to targets units with non-significant FCF (t-test, *** indicates *p <* 10^*−*6^).

We hypothesized that, when manipulating source unit *j*, its effect on target unit *k* could be predicted by the FCF estimated in the absence of perturbation (Fig. 4D). Specifically, we tested whether stimulation of source unit *j* would exert effects only on those target units *k* that have significant FCFs, *F*_*kj*_, but no effects on units whose FCFs were not significant. We found that the perturbation effects on the target units were localized to units with significant FCFs (dots in Fig. 4D; Fig. 4E). No effects were detected on pairs with non-significant FCF. In particular, we found that pairs where the stimulated source was in Y and the target in X did not show any significant effects of perturbations (Fig. 4E); this was expected given the absence of feedback couplings *Y→X*. Conversely, we found that source-target pairs with significant interventional connectivity after perturbation had much larger resting state FCF compared to pairs with non-significant interventional connectivity (Fig. 4F). Two crucial features of the FCF, underlying its predictive power, were its directed structure and its causal properties.

We thus conclude that the causal effect of perturbations on network units can be reliably and robustly predicted by the FCF inferred during the resting periods (i.e., in the absence of the perturbation).

### E. Causal flow from sparse recordings in spiking circuits

To apply our predictive framework to cortical circuits in behaving animals, we aimed at extending the method outlined above to encompass the following additional features of real neural circuits. First, while the rate network of Figs. 2-4 comprised real-value continuous rate units, neurons in cortical circuits exhibit spiking activity. Second, while the rate network was based on anatomical weights of either sign for a given source and weak recurrent couplings, cortical circuits are characterized by strong recurrent connectivity [10–13] obeying Dale’s law, with cell-type specific connectivity of excitatory (E) and inhibitory (I) neurons. Third, while so far we tested robustness of FCF inference to external shared and private gaussian noise, population activity in awake behaving animals exhibits super-Poisson variability, including both variability in firing rate fluctuations and spike timing [30, 43–45]. Fourth, commonly-used multi-channel electrode arrays typically yield sparse recordings of the underlying circuit activity: the number of active contacts in these electrodes (tens to hundreds) captures activity from only a small fraction of neurons in a circuit (typically *<* 1%). Crucially, each electrode records the aggregate spiking activity of a neural cluster, namely, a small number of neurons in a cortical column surrounding the electrode, some of which can be isolated as single units. We thus sought to extend our methods to address these critical issues, by performing a series of simulated experiments on a spiking neural network. Can we reliably infer functional causal flow between recorded neurons using sparse recordings of spiking activity at the level of neural clusters?

We inferred the FCF from sparse recordings of spiking activity in a large simulated cortical circuit (Fig. 5). In this model E and I spiking neurons were arranged in clusters, consistent with experiments supporting the existence of functional clusters in cortex [46–49]. In the model, E/I pairs of neurons belonging to the same cluster have potentiated synaptic couplings, compared to weaker couplings between pairs of neurons belonging to different clusters. Resting period activity in the clustered network displayed rich spatiotemporal dynamics in the absence of external stimulation, whereby different subsets of clusters activated at different times (with a typical cluster activation lifetime of a few hundred ms, Fig. 5A-B). These metastable dynamics were previously shown to capture physiological properties of resting activity in cortical circuits, in particular, the large variability in firing rate and spiking activity observed in awake behaving animals [28, 30–32, 50, 51].

**FIG. 5.**
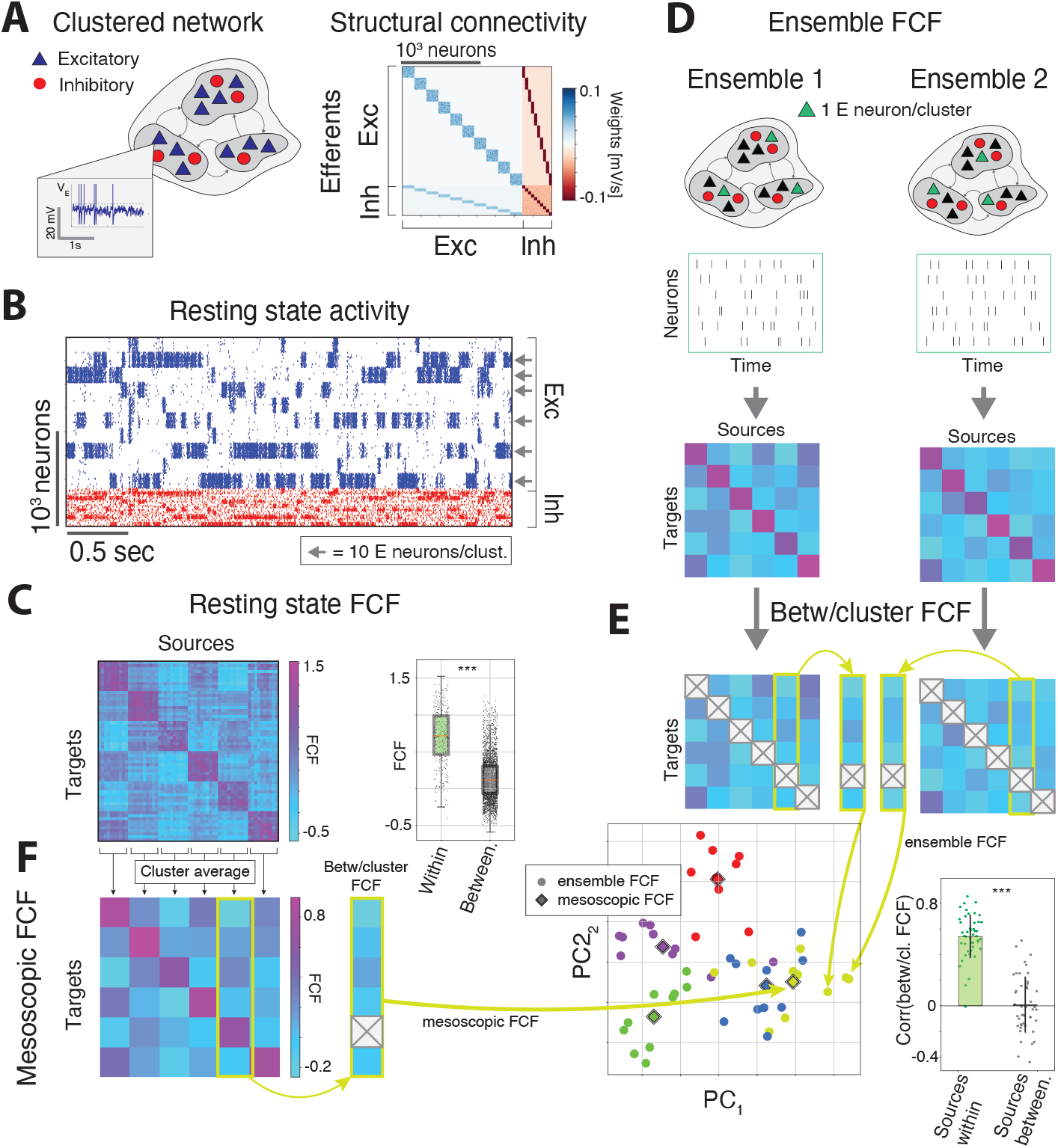
Mesoscopic causal flow in a sparsely recorded spiking circuit. A) Left: Schematics of the spiking network architecture with E and I neural clusters, defined by potentiated within-assembly synaptic weights. Inset shows membrane potential trace from a representative E cell. Right: Structural connectivity of the network. Blue and red represent positive and negative weights, respectively. B) Raster plots from a representative trial showing the entire network activity during the resting periods (blue and red marks represent action potentials of E and I cells, respectively; neurons arranged according to cluster membership). C. Left: Functional causal flow (FCF) between sparsely recorded activity (10 E cells per cluster were recorded, only six clusters firing above 5 spks/s on average were recorded; grey arrows in panel B). Right: FCF reveals the separation between pairs of neurons belonging to the same cluster (green) and different clusters (orange), reflecting the underlying anatomical connectivity (mean*±*SD, t-test, ***= *p <* 10^*−*20^). Data points represent neuron pairs. D) Ensemble FCFs inferred from two different recorded ensembles (one E cell were recorded from each of six clusters). E) Top: The between-cluster causal vectors (i.e., the columns of the off-diagonal ensemble FCF matrices) reveal the existence of a cluster-wise structure and grouping according to the cluster membership of their source neurons. Bottom left: Principal Component Analysis of between-cluster causal vectors, color coded for different clusters. Circles and rhomboids represent, respectively, between-cluster ensemble and mesoscopic vectors, the latter obtained from the mesoscopic FCF in panel F. Bottom right: Pearson correlations between vectors whose sources belong to the same cluster, left, or different clusters, right (t-test, *** indicates *p <* 10^*−*20^). F) Mesoscopic FCF based on the aggregated activity within clusters.

We estimated the FCF from short periods of resting activity (8 simulated seconds) from sparse recordings of E neurons (Fig. 5C, 10 neurons per cluster, 3% of total neurons in the network, only neurons with firing rates above 5 sp/s on average were retained for analyses). Visual inspection of the sparse FCF in Fig. 5C suggested the presence of two causal functional hierarchies in the circuit: the first hierarchy representing strong intra-cluster functional couplings and the second hierarchy representing weaker couplings between pairs of neurons in different clusters. The distributions of FCFs for pairs of neurons belonging to the same cluster, was significantly larger than the distribution of FCFs for the pairs belonging to different clusters (Fig. 5C, t-test, *p <* 10^*−*20^).

In primate recordings from multi-electrode arrays, each electrode typically records the aggregated spiking activity of a neural cluster surrounding the electrode. To model this scenario, we then examined the properties of the FCF between neural clusters in our simulated network. We investigated whether any hierarchy was present in the causal functional connectivity between different clusters. Close inspection of the FCF for pairs of neurons belonging to different clusters revealed the existence of clear off-diagonal blocks, suggesting the presence of a structure in the FCF between different clusters (Fig. 5C) at the *mesoscopic* level, namely, at the level of neural populations rather than single neurons.

We sought to quantify this mesoscopic structure by testing whether FCF between pairs of neurons, *F*_*ij*_ where *i ∈A* and *j ∈B*, encoded the identity of the target cluster *A* and source cluster *B* that the neurons were sampled from. We thus sampled small “ensemble FCF” matrices, obtained from subgroups of neurons, each consisting of one randomly sampled excitatory neuron per cluster (Fig. 5D, six neural clusters were considered). We hypothesized that, if the ensemble FCF captured the identity of the clusters from which neurons were recorded, then the causal vectors 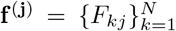 corresponding to source neurons *j* in the same cluster would be highly correlated, thereby allowing us to infer the network connectivity from sparsely recorded neurons Fig. 5D demonstrates this point for *N* = 6 clusters.

For this analysis, diagonal elements of FCF matrix should be removed. A naive dimensionality reduction on the columns of the full FCF matrix would have trivially led to the emergence of groups simply due to the fact that the diagonal entries of the FCF (self-reconstructability of the sources) were much larger than the off-diagonal ones *F*_*ii*_ *>> F*_*ij*_ |_*j/*=*i*_. This fact reflected the strong hierarchy in the intra- vs. inter-cluster FCF discussed above because we had sampled one neuron per cluster. In order to control for this diagonal effect and examine the smaller inter-cluster structure effects, we removed the diagonal elements from the *N×N* ensemble FCF matrices, and considered *N−*1-dimensional “between-cluster” causal vectors 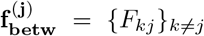 (Fig. 5E). We found that the between-cluster causal vectors grouped in Principal Component space according to the cluster membership of their sources (Fig. 5E). We further confirmed the existence of this mesoscopic structure by showing that the correlation of between-cluster causal vectors from sources belonging to the same cluster was much larger than those obtained from sources in different clusters (from Fig. 5E). These results show that FCF is a property shared by all neurons within the same neural cluster.

Building on this insight, we thus introduced the *mesoscopic* FCF as the coarse-grained causal flow between neural clusters, defined as the average FCF of the neurons in that cluster (block-average of FCF, see Fig. 5F).

Between-cluster causal vectors from the mesoscopic FCF stand at the center of each group of causal vectors from the between-cluster ensemble FCF (Fig. 5E), thus recapitulating the causal properties of each neural cluster. This mesoscopic causal flow is an emergent property of the clustered network dynamics and arises from the only source of quenched heterogeneity in the network, namely, the Erdős-Rényi sparse connectivity in the structural couplings. The strength of the underlying anatomical couplings for pairs of neurons belonging to two different clusters A and B depends only on the identity of the two clusters (see Methods). Namely, any pairs of neurons from clusters A and B share the same coupling strength, which in turn is different from the coupling between pairs of neurons belonging to clusters A and C. Thus the anatomical connectivity is a mesoscopic property of neural cluster pairs, and it is reliably captured by the mesoscopic FCF.

Together, these results uncovered a hierarchy of causal flow in a biologically plausible model of recurrent cortical circuits with functional clusters. The hierarchy separates large within-cluster FCF from smaller between-cluster FCF. Moreover, we found that the mesoscopic FCF revealed the functional organization between neural clusters. Crucially, the mesoscopic FCF suggests that our theory can be reliably applied to aggregate spiking activity of neural clusters recorded with micro-electrode arrays.

### F. Predicting perturbation effects from causal flow in spiking circuits

Is resting state FCF predictive of perturbation effects in the case of a sparsely recorded spiking network? To investigate this question, we devised a stimulation protocol whereby we briefly stimulated single neural clusters and examined the effect of stimulation on the activity of target ensemble neurons (Fig. 6). This stimulation protocol was designed to model perturbation experiments in alert monkeys (see below), where a brief electrical microstimulation of a single electrode on a multi-electrode array directly perturbs the cortical column surrounding the electrode, represented in our model by a neural cluster. Perturbation effects in the model were estimated by comparing the network activity immediately following the offset and preceding the onset of the stimulation (see Methods and Fig. 6A, representative clusters 2 and 3 were stimulated). The effects of stimulating a specific source neural cluster were encoded in the perturbation vector 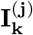, estimated for each source neuron *j* in the stimulated cluster, and for the sparsely recorded target neurons *k*.

**FIG. 6.**
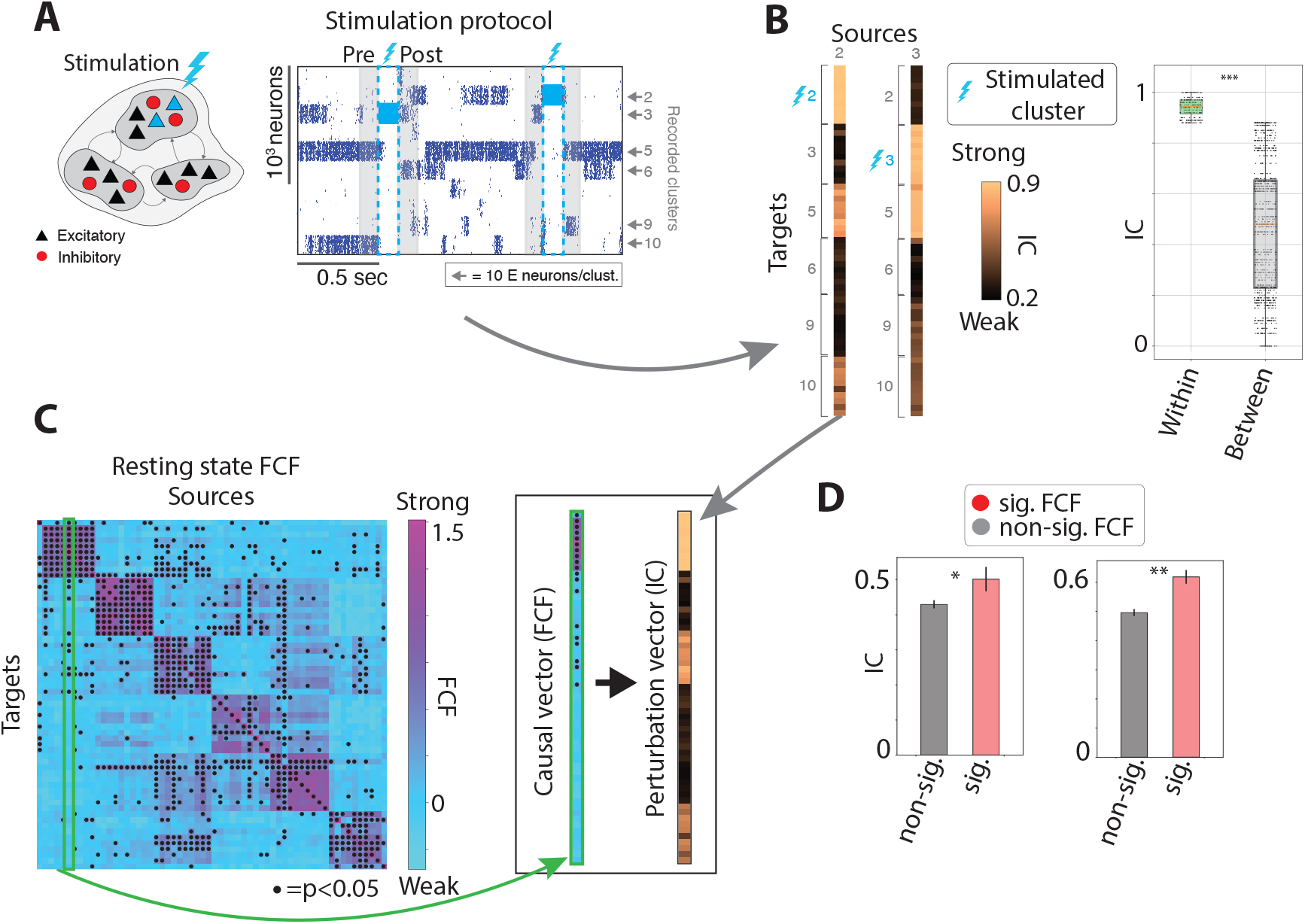
Functional causal flow predicts perturbation effects in spiking circuits. A) Perturbation protocol in the spiking network model of Fig 6. In this example, cluster 3 is stimulated followed by stimulation of cluster 2 (100ms blue interval in the raster plot). B) We used interventional connectivity to estimate the magnitude of response perturbations for a subset of target cells in clusters 2, 3, 5, 6, 9, and 10. ICs were calculated based on the spike count distributions during post-vs. pre-stimulation intervals (shaded gray areas; only clusters with average firing rate above 5 sp/s were considered; 10 E neurons per recorded cluster). Perturbation effects were larger for target neurons in the same cluster as the stimulated one (Within), compared to targets in different clusters (Between) (t-test, *** indicates *p <* 10^*−*20^). C) Between-cluster causal vectors obtained from resting activity FCF were used to predict between-cluster perturbation vectors (black dots indicate significant FCF pairs, *p <* 0.05). D. Resting state FCF predicts perturbation effects of stimulating source neurons in clusters 2 (left) and 3 (right). Red and gray bars show average IC for target units with significant and non-significant FCF for each source. * and ** indicate *p <* 0.05 and *p <* 0.01, respectively. Error bars are s.e.m.

We found that perturbations exerted a source-specific effect on the network activity, whereby stimulating neurons in different clusters led to differential patterns of responses across the clusters (Fig. 6B). We tested whether FCF was predictive of these perturbation effects. We had found above that the FCF between pairs of neurons belonging to the same cluster was much stronger than between pairs belonging to different clusters (Fig. 5C). We found the same hierarchy in the perturbation vectors, whereby the perturbation effects for targets belonging to the same stimulated cluster were much larger than for targets in different clusters (Fig. 6B). This hierarchy of perturbation effects closely matched the hierarchy present in the structural connectivity (Fig. 5A) and found in the resting state FCF (Fig. 5C). Overall, we found a strong correlation between resting state FCF and perturbation effects (Fig. 6D).

A crucial feature of the FCF was that it captured the mesoscopic structure of causal flow between different clusters (between-cluster FCF, Fig. 5). We thus set out to test whether between-cluster FCF could predict response perturbations of target neurons belonging to clusters different from the stimulated one. For this analysis, we consider “between-cluster” causal vectors and perturbation vectors obtained by masking all targets belonging to the stimulated cluster (Fig. 6C). We tested the predictive relation between resting state FCF and perturbation vectors, by separating the target neurons into those with significant FCF (“functionally connected”) and non-significant FCF (“unconnected”) for each stimulated source. Our theory predicted that perturbation effects would be stronger for functionally connected targets. Fig. 6D confirms this prediction, thus demonstrating that causal flow is predictive of perturbation effects even in the sparse recordings regime (Fig. 6D).

Overall, these results show that in a biologically plausible cortical circuit model with functional clusters, the effects of perturbations can be robustly captured at the mesoscopic level of neural clusters even using a sparse recording regime, typical in real experiments, which we investigate next.

### G. Inferring functional causal flow in the prefrontal cortex of alert monkeys during resting periods

To test our theory, we performed recording of spiking activity and microstimulation in the monkey prefrontal cortex (pre-arcuate gyrus, area 8Ar) during a period of quiet wakefulness (resting) while the animals were sitting awake in the dark. The experiment had two phases (Figs. 1 and 7). In the first phase, we recorded population neural activity from a multi-electrode array (96-channel Utah array, with roughly one electrode in each cortical column in a 4*×*4mm^2^ area of the cortex), estimating the FCF between pairs of neural clusters (multiunit activities collected by each recording electrode). In the second phase, we perturbed cortical responses by delivering a train of biphasic microstimulating pulses (15 *μA*, 200 Hz) to one of the clusters for a brief period (120ms), recording population neural activity across the array before and after the stimulation.

**FIG. 7.**
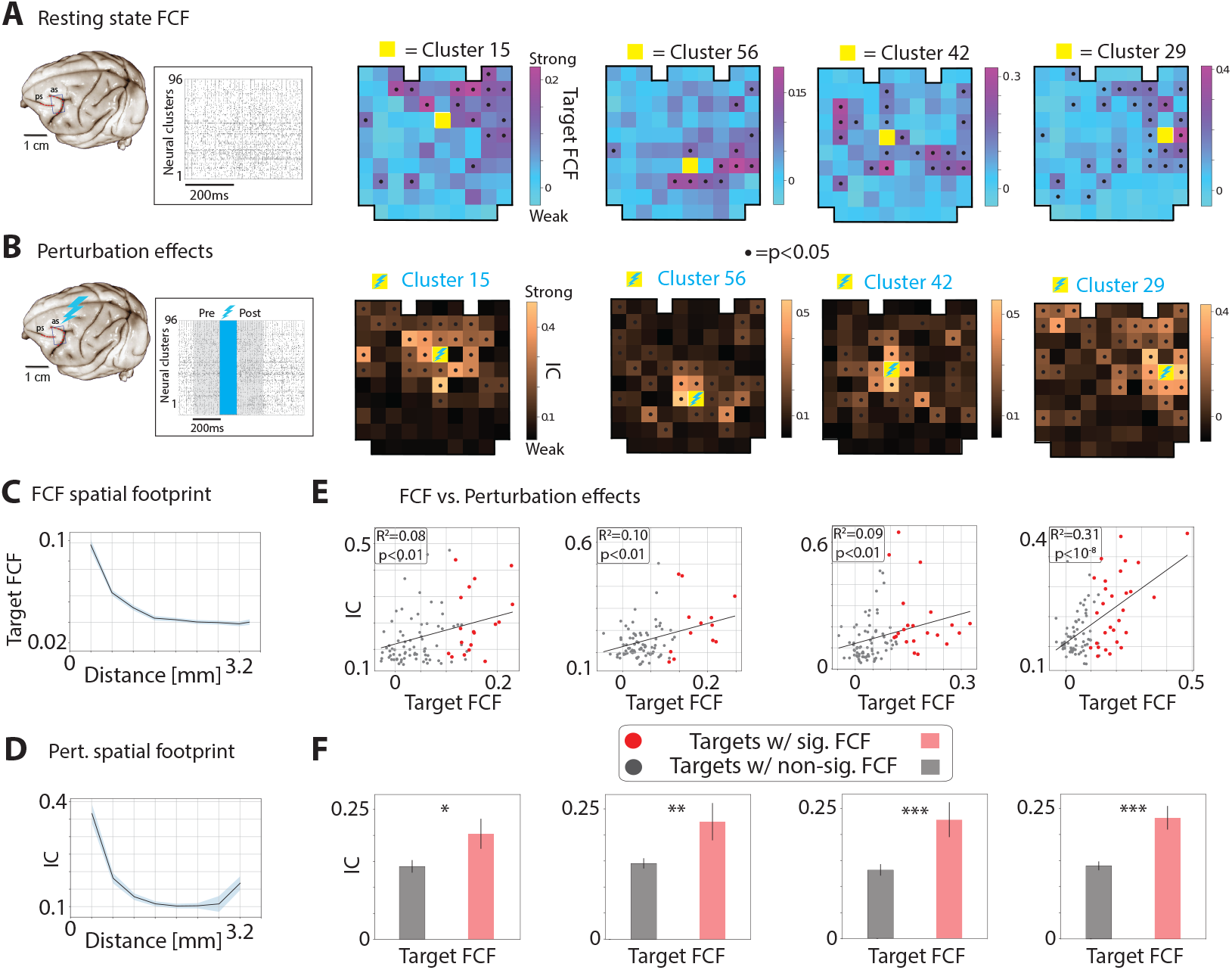
Functional causal flow predicts perturbation effects in the prefrontal cortex of resting, alert monkeys. A) Left: Ensemble spiking activity of pre-arcuate gyrus recorded with a 96-channel multi-electrode array. Black tick marks are spikes from each neural cluster, defined as the aggregated spiking activity around each recording electrode. Right: Causal vectors for four representative source clusters. Yellow squares show the location of the source cluster on the array. FCF causal vectors for each source are also overlaid on the array geometry (black dots represent significant FCF values, established by comparison with surrogate datasets, *p <* 0.05, see Fig. S2 and Methods). For full FCF matrix see Fig. S3. B) Left: Response perturbations caused by microstimulation of electrode 15 (120ms biphasic 15*μ*A stimulation train, blue shaded area) across clusters. Right: Interventional connectivity for the four stimulated clusters measured based on spike count distributions in 200ms intervals preceding and following the stimulation (gray shading in the population raster plot). Black dots represent significant ICs (*p <* 0.05). C) FCF decays with increasing distance of the target and source clusters. D) Perturbation effects for the four stimulated clusters decays for more distant target clusters. Shading indicates s.e.m. in panels C and D. E) Resting state FCF predicts perturbation effects. For each stimulated source, target clusters with larger FCF are more strongly impacted by microstimulation, indicated by larger ICs. Gray and red data points represent targets with non-significant and significant FCF, respectively (*p <* 0.05). Black lines are linear regressions. *R*^2^ and slope p-value are reported. F) For each stimulated source in panel E, aggregated perturbation effects are larger over targets with significant FCF vs. targets with non-significant FCF. Error bars are s.e.m. *,**,*** indicate *p <* 0.05, 0.01, 0.001 (t-test).

We first examined whether functional causal flow could be estimated reliably for the recorded population, which constituted a small fraction of neurons in the circuit. Our modeling study suggested that FCF can be defined at the mesoscopic level as a property of functional assemblies or neural clusters: FCF inferred from different neurons recorded from the same cluster yielded consistent results (Fig. 5). Following previous experimental evidence supporting the existence of assemblies in monkey pre-arcuate gyrus [49], we reasoned that the recorded neurons around each electrode may represent sparse samples from a local cortical cluster. This experimental setup thus provided a similar scenario to the one validated in our modeling study (Fig. 5).

We found that FCF was characterized by a complex set of spatiotemporal features (Fig. 7A, for the full 96 *×* 96 FCF matrix see Fig. S3, see Methods for details and for model selection). In Fig. 7A we show four representative 96-dimensional causal vectors representing the FCF for four different source clusters (channels 14 and 56 from session 1 and channels 42 and 29 from session 2; these source clusters were microstimulated in the second phase of those sessions, see the following section). We overlaid the causal vectors onto the array geometry (location of recording electrodes in the array) for illustration (each overlaid causal vector in Fig. 7A corresponds to a specific column of the full FCF matrix in Fig. S3).

Comparison of the causal vectors across source clusters revealed remarkable features about the structure of the functional connectivity. First, FCF is channel-specific, namely, it depends on the source clusters whose activity is being reconstructed. Second, each causal vector shows a hierarchical structure, with significant FCF rkfor a subset of targets, while most targets cannot reconstruct the source activity (Fig. 7A). This result is qualitatively consistent with the FCF obtained from our models (Figs. 3 and 6), supporting the hypothesis of functional hierarchies embedded within prefrontal cortical circuits [49].

Is the FCF of a source cluster to different target clusters uniformly distributed across the array, or is there a preferential spatial footprint of FCF? We found a spatial gradient whereby FCF was largest in the target clusters immediately surrounding the source cluster, while FCF for distant targets typically plateaued at low but nonzero values (Fig. 7C).

We thus conclude that FCF inferred during the resting periods is cluster-specific and reveals a hierarchy of functional connectivity where functionally downstream neural clusters are spatially localized around the source cluster. These results extend previous correlation analyses of spatial clusters in alert monkeys [49] highlighting a spatial gradient of directed functional couplings at the mesoscale level.

### H. Perturbation effects on cortical circuits in alert monkeys

We next proceeded to examine the effect of microstimulation on the cortical activity in alert monkeys. We estimated perturbation effects using interventional connectivity, which uses Kolmogorov-Smirnov test statistics to calculate the dissimilarity of the activity distribution of neural clusters in the intervals preceding the onset and following the offset of the stimulation for each pair of stimulated source and recorded target clusters (see Fig. 7B). We focused on the activity after offset as opposed to during the stimulation period to minimize the effect of stimulation artifacts on the recording apparatus.

We first examined the spatiotemporal features of stimulation effects. We found that perturbations exerted a strong effect on ensemble activity, and that these effects where specific to which electrode was stimulated (Fig. 7B; perturbation effects for each stimulated source *j* are visualized as a perturbation vector **I**^(**j**)^ overlaid on the array geometry). By comparing the effects of stimulating a single electrode on all target clusters, we found a hierarchical structure with strong effects elicited in specific subsets of target clusters. The identity of strongly modulated targets was specific to the stimulating electrode. We found a spatial gradient in perturbation effects, whereby distant targets were less affected by perturbation, though the effects were nonzero even far away from the stimulated electrode (Fig. 7D).

### I. Predicting perturbation effects from resting activity in alert monkeys

Our theory posits that the effects of stimulation of source cluster *j* on the target neural clusters can be predicted by the corresponding causal vector **f** ^(**j**)^ inferred at rest (i.e., a column of the FCF matrix). Specifically, we predict that perturbing a source cluster exerts a stronger effect on those target clusters which have a stronger functional connectivity to the source as defined by the source causal vector. Visual inspection of the resting state FCF causal vectors (Fig. 7A) and the map of perturbation effects (Fig. 7B, perturbation vectors) suggests strong similarites for any given source. We confirmed this quantitatively using Pearson correlations between causal vectors and perturbation vectors (Fig. 7E). Additionally, the effect of stimulation was significantly stronger on targets with stronger functional connectivity to the stimulated cluster compared to targets with weak functional connectivity, as predicted by our theory (Fig. 7F). The predictive power of FCF held at the level of single stimulated sources, thus achieving a high level of granularity in prediction.

Because both the FCF and perturbations displayed a characteristic decay proportional to the distance from the source electrode, we tested whether the predictive relationship between them still held after controlling for spatial distance. After removing this spatial dependence, the predictive relation between FCF and perturbation effects still held for the residuals (Fig. S4), thus confirming that the FCF predicts perturbation effects above and beyond what is expected from a spatial decay away from the source cluster. These results demonstrate that the functional causal flow estimated from sparse recordings during the resting periods accurately predicts the effects of perturbation on the neural ensemble, thus establishing the validity of our theory in cortical circuits of alert monkeys.

### J. FCF predictive power outperforms other causality methods

The approach to measuring causality using FCF overcomes some of the well-known limitations of traditional causality measures based on information transfer, such as Granger causality (GC, [52]), as well as its direct generalization known as transfer entropy [53]. We refer the reader to the Supplementary Section A for a detailed examination of the technical and conceptual differences between FCF and these alternative methods.

Here, we compare the performance of FCF in predicting perturbation effects with alternative methods based on information theory, specifically Granger causality in its univariate (GC) and multivariate (MGC) formulations; nonlinear GC (NGC) with radial basis functions for autoregression; and transfer entropy (TE). We first used each of these methods to estimate directed functional connectivity based on resting periods in the simulated continuous rate network (Fig. S6). We then examined the correlation between each causality index and the effects of perturbations, the latter estimated as the interventional connectivity (Fig. 4). For a causality index to successfully predict the effects of perturbation, we expect a large positive correlation between a source-target pair’s IC and the value of the causality index.

We found that FCF had a significantly larger correlation with IC (*r* = 0.50, *p* = 1.7 *×* 10^*−*6^) compared to all other indices (Fig. S6). Among the other indices, GC had a significant positive correlation as well (*r* = 0.32, *p* = 2.8*×*10^*−*3^), but TE and NGC had no significant correlations. These results were corroborated by an additional test that showed source-target pairs with significant IC had larger FCF (*p* = 1.8*×*10^*−*9^) and GC (*p* = 9 *×* 10^*−*5^) than pairs with non-significant IC, whereas the other indices did not exhibit such a difference. From these comparative analyses on simulated data we concluded that, among the causality indices, FCF performs best. GC is also predictive of perturbation effects, although to a lesser degree than FCF, while NGC, TE, and MGC do not exhibit predictive features.

We then deployed the causality indices to infer the causal structure of the multielectrode recordings from the monkey prefrontal cortex (Fig. S7). As in the comparative analysis of the simulated rate network above, we estimated the correlation between causality indices and IC, and the difference in causality indices for source-target pairs with significant vs. non-significant values of IC. While the spatiotemporal profile of FCF closely resembled that of the IC, this similarity was less pronounced for the other causality indices. We found a large positive correlation for FCF (significant in both monkeys). GC showed a positive, albeit smaller, correlation as well (significant in only one of the two monkeys). The other indices showed no consistent predictive power.

We thus conclude that among the alternative causality indices, the only one capable of predicting perturbation effects to some degree is GC, which shows decent performance in the simulated rate network and a positive trend in the experimental data; albeit significantly worse than FCF in all cases. The other indices NGC, MGC, TE fail to predict perturbation effects. This comparative analysis thus highlights the advantages of FCF in predicting the effects of perturbations from resting data in alert monkeys.

## III. DISCUSSION

Predicting the effect of targeted manipulations on the activity of cortical circuits is a daunting task but it could be achieved by capturing the causal functional interactions in the circuit. A central challenge using common multi-electrode arrays in monkeys and humans is the extremely sparse recording regime, where the activity of only a small fraction of neurons in a circuit is observed. In this regime, traditional methods fail due to unobserved neurons and common inputs to the circuit.

Here, we demonstrated a new framework for predicting the effect of perturbations to a cortical circuit based solely on the causal interactions within a circuit inferred from sparsely recorded spiking activity at rest. We validated the method on ground truth data showing that functional causal flow (FCF) captures the functional connectivity in a biologically plausible model of a cortical circuit, even when recording only a small fraction of the network’s neurons. The FCF inferred at rest predicts the effects of perturbations encoded in the network interventional connectivity matrix. FCF is remarkably robust to external noise and common inputs. Using FCF inferred during the resting periods in the network, we predicted the effect of perturbing a neural cluster on the rest of the recorded neurons, revealing the set of target neurons with directed functional couplings to a given source. Remarkably, when applying FCF to spiking activity from ensemble recordings in alert monkeys, we were able to reconstruct the causal functional connectivity between the recording electrodes using resting activity. The resting state FCF predicted the effect of single-electrode microstimulation on target electrodes which were classified as functionally coupled to the source. A detailed comparison of our methods to Granger causality and other alternative methods showed that FCF outperforms GC on simulated and real data, and, in particular, GC fails when applied to the latter. Our results establish a new avenue for predicting the effect of stimulation on a neural circuit solely based on sparse recordings of its resting activity. They also provide a new framework for discovering the rules that enable generalization of resting state causal interactions to more complex behavioral states, paving the way toward targeted circuit manipulations in future brain-machine interfaces.

### A. The role of resting activity

In traditional neurophysiological studies, resting activity is defined as a pre-stimulus background activity, immediately preceding stimulus presentation, and it is regarded as random noise or baseline, devoid of useful information [54–56]. However, recent results have challenged this widely held picture, producing evidence that resting activity may encode fundamental information regarding the functional architecture of neural circuits.

Studies investigating the dependence of neural responses on the background activity, quantified with local field potentials [57, 58], single neuron membrane potentials [59], or population spiking activity [42], found that it encodes information about the animal’s behavioral state, including even fine grained movements [60–62]. Moreover, resting activity immediately preceding stimulus onset predicts the trial-to-trial variability in stimulus evoked response, potentially explaining the observed dependence of sensory responses on the underlying state of the network [63, 64].

Recent studies have suggested that resting activity is finely structured, containing information on the functional architecture of the neural circuits [49, 65, 66] and providing a repertoire of network patterns of activation [30, 67, 68], potentially linked to developmental plasticity [69, 70]. The population coupling of single neurons estimated during the resting periods *in vivo* is correlated with the synaptic input connection probability measured *in vitro* from the same cortical circuit [71]. Also, in neuronal cultures, the causal functional connectivity inferred from ongoing activity is predictive of the structural connectivity estimated from electrical stimulation: functionally downstream neurons have faster response latency to stimulation compared to functionally upstream neurons [26].

### B. Circuit models of resting activity

We validated our theoretical framework for causal inference using two classes of models: a continuous rate network and a spiking network model of resting activity in a cortical circuit. The latter is a biologically plausible model based on a recurrent spiking network where excitatory and inhibitory neurons were arranged in functional assemblies [28, 30, 32, 51]. Experimental evidence including multielectrode recordings in behaving monkeys [49] strongly supports the existence of functional assemblies in cortex [46–48, 72]. Clustered spiking networks capture complex physiological properties of cortical dynamics during resting and stimulus-evoked activity due to the metastable dynamics of cluster activations. Such physiological properties include context- and state-dependent changes in neural activity and variability [28–30, 50, 51, 73]; as well as neural correlates of behavior and cognitive function such as expectation, arousal, locomotion, and attention [31, 32, 51].

### C. Estimating functional connectivity

Estimating functional connectivity is a formidable task, especially susceptible to errors in the presence of strong recurrent couplings, noise, and unobserved common inputs ubiquitous in cortical circuits. Even when unlimited data are available, sophisticated methods typically fail in the presence of strong correlations between unconnected neurons [74].

We have defined functional interactions as the causal interaction of cortical neurons. Existing methods for estimating functional interactions between multidimensional time series include linear regression [75], Granger causality (GC) [76], and inter-areal coherence [77, 78]. While correlation-based methods are problematic for weak correlations, entropy-based methods such as transfer entropy [79] are extremely data hungry. Detecting a clear causal relationship by transfer entropy [79] or Granger causality [16–18, 80] is not straightforward unless the system’s dynamical properties are well known. Further, confounding effects of phase delay [22], self-predictability in deterministic dynamics [21] or common inputs [81, 82] limit the usefulness of Granger causality and transfer entropy. Alternatives such as inverse methods based on Ising models utilize time-consuming learning schemes [83] though recently faster algorithms have been proposed [14, 84]. Other approaches applicable to spike trains include generalized linear models [85] or spike train cross-correlograms [86]. Remarkably, the latter method was successfully validated using optogenetic perturbations *in vivo*.

Inferring causal functional connectivity from *extremely sparse recordings* of neural activity is a long standing problem. Our method relies on delay embedding techniques used for reconstructing nonlinear dynamical systems from their time series data with convergent crossmapping [21]. Crucially, convergent cross-mapping was designed to work precisely in the sparse recording regime [23, 24], where other methods fail. While this powerful framework has been successfully applied in ecology [21], and previously applied to ECoG data [27] and in neuronal cultures [26], here we pioneered its use for estimating causal functional connectivity from spiking activity of a neural population in awake monkeys. Using synthetic ground truth data from recurrent spiking networks, we showed that FCF can be reliably estimated using extremely sparse recordings and very short samples of neural activity (tens of seconds, Fig. 5).

We performed a detailed comparison of FCF to univariate, multivariate and nonlinear Granger causality, and transfer entropy. We found that FCF outperformed all other methods. Among the latter, only univariate GC was predictive of perturbation effects, although it underperformed FCF both regarding the resting state inference of functional connectivity, as well as the ability to predict perturbation effects. The decent performance of GC, compared to the other alternaive methods, might be due to the fact that GC is relatively less data hungry.

### D. Hierarchical structures in cortical circuits

Previous studies strongly support the existence of a hierarchical structure in brain architecture both at the whole-brain level [87, 88] as well as locally within single cortical areas [63, 71]. In the latter case, ensemble neurons were ranked based on the degree to which their activity correlated with the average population activity (i.e., soloist and choristers). Here, we took a step beyond correlational analysis and revealed the hierarchical structure of causal interactions using resting period data. For each source channels, we were able to classify its targets as functionally “upstream” or “downstream” within the network’s causal flow. When the ground truth structural connectivity is known, as in our simulated networks, we showed that the causal hierarchy may reflect structural couplings and the existence of neural assemblies. Surprisingly, we found that causal hierarchies within a local circuit may naturally emerge from heterogeneities in recurrent couplings between neural assemblies, even in the absence of a feedforward structure.

In cases where the structural connectivity is not accessible, as in alert primate recordings, we showed that the causal hierarchy inferred from resting periods predicts the effect of perturbations via electrical microstimulation. Specifically, perturbation of a source electrode affects more strongly target electrodes which are functionally coupled to the source compared to those that are not functionally connected. Although both FCF and perturbation effects exhibited a spatial decay with the distance from the source channel (consistent with previous results on spatial correlations in primate cortex [49, 89]), we found that the FCF predicted perturbation effects above and beyond what expected from the common spatial decay (Fig. 7G). The spatial decay profile of our microstimulation protocol reached a minimum 2mm away from the stimulation site. We thus found a much larger influence radius compared to optogenetic stimulations of single neurons in mouse visual cortex, where the perturbation effects was confined to a radius of about 25*μ*m [90]. This suggests that our perturbation, although quite weak in magnitude and duration and below perceptual threshold, likely recruited larger neural ensembles.

### E. Microstimulation effects on neural activity in primates

Microstimulation experiments have played a crucial role for our understanding of the organization and function of neural circuits in the primate brain. Among many successful examples are microstimulation of motion-selective middle temporal (MT) neurons to alter choice [1], reaction time [91], or confidence [92] of monkeys performing a direction discrimination task. However, outside of sensory or motor bottlenecks of the brain, the use of microstimulation (or other perturbation techniques) is fraught with challenges. In many regions of the primate association cortex, neurons have complex and task-dependent selectivities. Identifying these selectivities is often time-consuming and may not be always possible. Further, perturbation of the activity of these neurons does not necessarily lead to behavioral changes commensurate with their empirically defined selectivities, partly because selectivities in one task condition may not gen-eralize to others and partly due to network effects of perturbations beyond the directly manipulated neurons.

### F. Model-based approach to perturbation experiments in primates

Our approach to quantify FCF based on the activity of a large neural population spread out in multiple neighboring cortical columns provides an easily implementable solution with many advantages. First, we directly assess the network effects of the activity of each neuron, and thereby generate predictions about the impact that perturbing the activity of one cluster of neurons will have on the rest of the population. Second, population activity has proven quite powerful in revealing the neural computations that underlie behavior, with features that are robust to the exact identity of the recorded neurons and their complex selectivities [93–98]. Previous studies characterized the effect of perturbations on single neurons in cortical circuits [90] and clarified the structure of anatomical circuit connectivity required to produce the observed effects [99]. In a similar fashion, we defined interventional connectivity as the effect of the perturbation of a single node on the activity of all other observed nodes. The novelty of our approach, compared to earlier studies, is the ability to predict the interventional connectivity from the FCF inferred at rest. We suggest that characterizing FCF during a task and using our model-based approach to predict the impact of a variety of perturbations on the population level representations offer an attractive alternative to the traditional trial-and-error approaches where different neural clusters are manipulated in search for a desirable behavioral effect. We speculate that our model-based approach may lead to crucial advances in brain-machine interfaces if one can use FCF inferred from resting activity to predict perturbation effects during a task – a direction we will actively pursue in the future. Previous studies in human surgical patients aimed at predicting the effects of intracortical stimulations from measures of functional connectivity. Changes in EEG frequencies induced by the stimulation at specific sites could be predicted from resting state EEG coherence [100]. The spatial location of stimulation-evoked potentials could be predicted from the resting state fMRI spectrum seeded at the stimulation site [101]. Alternative approaches aimed at predicting the effects of stimulation using network control theory, although at the whole brain level [102].

Two key challenges in interpretation of typical microstimulation experiments are: (i) indirect activation of distant neurons through the activation of the neural cluster around the stimulating electrode, and (ii) effects on fibers of passage that could cause direct activation of neurons distant to the stimulating electrode [103]. Our approach directly addresses the first challenge by mapping the FCF based on the ensemble activity. The second effect acts as noise in our approach because the FCF is quantified based only on the activity of the neurons recorded by the electrodes. The success of our approach (Fig. 7) suggests that this noise is not overwhelming. The robustness of our approach likely stems from the synergy of our delay embedding methods with our focus on the population neural responses and the large number of simultaneously recorded neural clusters in our experiments, which effectively capture key features of the intrinsic connectivity in the circuit (Fig. 2) [28, 30, 31, 49, 51]. A recent theoretical study [99] supports the viability of using biologically plausible models of cortical circuits in terms of E-I networks with structured connectivity to explain perturbation experiments in awake animals [90], although they do not attempt to predict the perturbation effects.

We carried out a detailed comparison of FCF with Granger causality (GC), which highlighted the conceptual and practical advantage of FCF. We found that GC performed reasonably well on simulated data from continuous rate network. Specifically, although GC was less effective at detecting the functional hierarchy present in the network, it allowed to predict the effect of perturbations using the functional connectivity estimated in the absence of perturbations. GC did not perform well on the clustered spiking network. Unlike FCF, GC failed when applied to the clustered spiking network, where it was not able to detect the hierarchical structure in the structural couplings, nor was it able to predict perturbation effects from resting state functional connectivity. We found a strikingly different performance between the two methods on ensemble spiking activity from awake monkeys as well. First, the functional connectivity map inferred from resting activity was notably different when using FCF or GC. Crucially, while FCF inferred at rest was predictive of the effects of electrical microstimulation, GC failed to do so. We expected both GC and FCF to perform well when the hierarchical network structure is clear and the dynamics can be approximated by a linear or piecewise linear structure, such as in the case of the Rossler attractor network. However, in the presence of strong recurrent couplings, complex nonlinear dynamics, common inputs, and sparse subsampling regime, such as in cortical ensemble activity, the assumptions underlying GC do not hold, leading to its expected failure to predict stimulation effects. On the other hand, FCF was designed to work precisely in the regime of cortical dynamics, and thus we expected it to perform well in all cases as confirmed by our results.

Current methods for establishing site efficacy for perturbation experiments are labor intensive, time consuming, and often unable to generalize beyond the limited task set they are optimized for. Here, we demonstrated a new statistical method capable of predicting the impacts and efficacy of a targeted microstimulation site using only the resting activity. Crucially, our method can directly be applied to monkeys and humans, where commonly used “large-scale” recording technologies often permit sampling from only a small fraction of neurons in a circuit (typically *<* 1%). Our method is thus likely to improve the safety and duration of the procedure, a key step toward targeted circuit manipulations for ameliorating cognitive dysfunction in the human brain, as well as development of future brain-machine interfaces.

### G. Implications of causal flow for neural coding in intervention experiments

Perturbations of cortical circuit activity have been prominently used to study neural computations and coding both *in vivo* [90, 104] and *in silico* [99]. We anticipate several avenues for our predictive framework to facilitate innovations on existing interventional approaches [105]. The first application is related to virtual sensation experiments, where an animal is trained to discriminate between artificial activity patterns induced by microstimulation, in the absence of sensory stimulation [106]. Because the FCF inferred at rest is predictive of microstimulation effects, one could test whether animals can more easily discriminate between artificial stimuli consisting of stimulating electrodes with high resting FCF, compared to electrodes with low FCF. This hypothetical experiment would test whether sensory and decisionmaking pathways are regulated by spontaneous activity patterns and functional connectivity.

The second direction is to intervene with microstimulation concomitant or following the presentation of sensory stimuli during delay-response tasks, with the goal of biasing the animal’s decision, such as in the classic MT microstimulation experiments in primates [1]. Intervention could be optimized by first estimating stimulus- and choice-selectivity during the first part of a recording session. Combining knowledge of FCF with selectivity, efficient intervention protocols could be designed by monitoring the animal’s decision in real time [93] during the delay period to rescue a wrong choice, by perturbing choice-selective electrodes with high FCF values.

More generally, the fact that FCF inferred at rest could recapitulate the effect of our microstimulations suggests that low-intensity perturbations may respect the natural correlational structure of population dynamics, representing so-called “in-manifold” perturbations [107]. Combining FCF-informed perturbation with knowledge of electrode selectivity could lead to designing perturbation experiments that do not disrupt the intrinsic neural manifold and minimize off-target effects [108]. This paradigm could help clarify the extent to which neural patterns in different areas are causally contributing to the animal’s deliberation during decision-making tasks and provide a new avenue for brain-machine interfaces to rescue cognitive deficits.

### H. Limitations of functional causal flow

The estimation of functional causal flow as we presented it in this study is based on pairwise comparison between time series. In order to move beyond neuron pairs, we introduced the mesoscopic causal flow between neural clusters, defined as the average FCF between pairs of neurons belonging to two clusters. This mesoscopic measure is also limited to pairs of clusters. In its current form, our method may not be directly generalized to detect triplets or higher order interactions. An interesting direction for future work is the generalization of our method to capture multi-neuron perturbations, namely, predicting the joint causal flow between two source neurons and one target neuron. This generalization could pave the way to predicting the effect of simultaneous multi-electrode perturbation on the activity of downstream neurons. A powerful alternative method for modeling triplet interactions is the Partial Information Decomposition [109]. We hope to return to this question in future work.

Another potential limitation of FCF stems from the fact that neuronal activity in frontal areas likely receives time-varying input from several other cortical and subcortical areas. Moreover, the recorded ensemble neurons may receive common inputs from unobserved neurons within the same local prefrontal circuit. Moreover, these sources of external input may change across different periods of the resting state, due to changes in the animal’s internal state such as arousal levels. These contextual effects might present a potential challenge when generalizing FCF predictions across different conditions (such as resting vs. task engaged sessions). Our results address this concern by showing that FCF estimates are robust to both private noise and time-varying external inputs, typically encountered in cortical circuits (Fig. 3). It is an interesting open question to estimate how FCF may generalize across different behavioral conditions and we hope to report on this in the future.

## ACKNOWLEDGMENTS

The authors would like to thank Rainer Engelken and Ramin Khajeh for advice on the simulations and Tim Gardner for discussions. AN was supported by NSF DBI-1707398 and by the Gatsby Charitable Foundation grant GAT3708. RK was supported by Simons Collaboration on the Global Brain (grant 542997), McKnight Scholar Award, Pew Scholarship in the Biomedical Sciences, and National Institute of Mental Health (R01 MH109180). LM was supported by National Institute of Neurological Disorders and Stroke grant R01-NS118461 (BRAIN Initiative) and by National Institute on Drug Abuse grant R01-DA055439 (CRCNS). SE was supported by NYU Dean’s Dissertation Fellowship Award. The authors declare no competing interests.

## Author contributions

L.M. and R.K. designed the project; L.M., R.K. and T.T. supervised the project; S.E. and R.K. collected the experimental data; A.N., F.F. and L.M. developed the models and analyses; A.N. ran model simulations and analyzed experimental data with help from F.F. and L.M.; A.N. wrote the open source software package; L.M., A.N., R.K. and F.F. wrote the manuscript.

## IV. MATERIALS AND METHODS

### A. Definitions

We define the **functional causal flow** (FCF) from unit *x*_*j*_ to *y*_*i*_ as the reconstruction accuracy *F*_*ij*_ of the activity of *x*_*j*_ given the activity of *y*_*i*_. The accuracy is quantified as the Fisher z-transform of the Pearson correlation between the *empirical* activity of unit *x*_*j*_ and its *predicted* activity obtained from unit *y*_*i*_, that is *F*_*ij*_ = *z*[*ρ*(*x*_*j*_|*y*_*i*_)]. The **source unit** *x*_*j*_ is the one whose activity is being reconstructed and the **target unit** *y*_*j*_ is the one used for the reconstruction.

The j-th **column** of the FCF matrix *F*_*ij*_ represents the reconstruction of the activity of a particular source unit *x*_*j*_, given the activity of *target* unit *y*_*i*_. The i-th **row** of the FCF matrix *F*_*ij*_ represents the degree to which the activity of a particular target unit *y*_*i*_ can reconstruct the activity of source units *x*_*j*_. A summary is provided in Table I.

### B. Network models

#### 1. Rate network

The network consists of 100+3 nodes, 3 of which belong to the subnetwork X following Rössler dynamics described by the equations below:

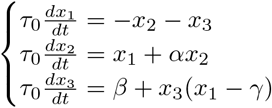

Other nodes in subnetwork Y evolve according the following dynamics:

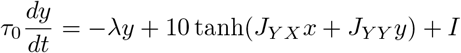

As observed from the above equations nodes in *X* are uni-directionally projecting to nodes in *Y*. The weight matrix *J*_*Y X*_ connecting *X* to *Y* is the product of a scalar *g*_*i*_ (connection strength) and a binary matrix where the elements are sampled from *Bernoulli*(*p*). The recurrent weight matrix *J*_*Y Y*_ is drawn from (0, *g*_*r*_). It is shown that increasing *g*_*r*_ will transition the network into a chaotic regime. See Table II for a description of the model parameters and their values. In our conventions for the rate network we set the unit of time to *τ* = 1ms.

**TABLE II.** Parameters for the rate network.

| Rate network simulations |  |  |
| --- | --- | --- |
| Parameter | Description | Value |
| $g_r$ | Strength of the recurrent weights | 4 |
| $g_i$ | Strength of $X$ to $Y$ weights | 0.1 |
| $p$ | Connection probability from $X$ to $Y$ | 1 |
| $\alpha$ | Rössler parameter | 0.2 |
| $\beta$ | Rössler parameter | 0.2 |
| $\gamma$ | Rössler parameter | 5.7 |
| $\lambda$ | Recurrent leak term | 1 |
| $\tau_0$ | Time constant | 1 ms |

#### 2. Spiking network

We modeled the cortical circuit as a network of *N* = 2000 excitatory (E) and inhibitory (I) spiking neurons (*n*_*E*_ = 80% and *n*_*I*_ = 20% relative fractions). Connectivity was Erdős-Renyi with connection probabilities given by *p*_*EE*_ = 0.2 and *p*_*EI*_ = *p*_*IE*_ = *p*_*II*_ = 0.5. When a synaptic weight from pre-synaptic neuron *j* to post-synaptic neuron *i* was nonzero, its value was set to 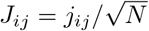, with *j*_*ij*_ sampled from a gaussian distribution with mean *j*_*αβ*_, for *α, β* = *E, I*, and variance *δ*. E and I neurons were arranged in *C* = 10 clusters of equal size. Pairs of neurons belonging to the same cluster had potentiated synaptic weights by a ratio factor 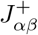, for *α, β* = *E, I*. Pairs of neurons belonging to the different cluster had depressed synaptic weights by a factor 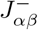. We chose the following scaling: 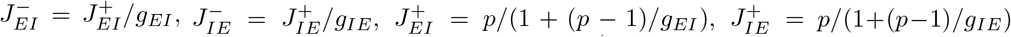 and 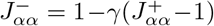 with *γ* = *f* (2 *− f* (*p* + 1))^*−*1^, where *f* = (1 *− n*_*bgr*_)*/p*. Network parameters were chosen to generate spontaneous metastable dynamics with a physiologically realistic cluster activation lifetime, consistent with previous studies [30–32, 110]. Parameter values are in Table III.

We used leaky-integrate-and-fire (LIF) neurons whose membrane potential *V* evolved according to the dynamical equation

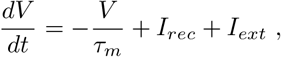

where *τ*_*m*_ is the membrane time. When *V* hits threshold 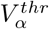 (for *α* = *E, I*), the neuron emits a spike and *V* is held at reset *V* ^*reset*^ for a refractory period *τ*_*refr*_. Thresholds were chosen so that the homogeneous network (i.e.,where all 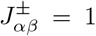) was in a balanced state with rates (*r*_*E*_, *r*_*I*_) = (2, 5) spks/s [31, 32, 111]. Input currents contained a contribution *I*_*rec*_ from the recurrent connections and an external current *I*_*ext*_ = *I*_0_ + *I*_*pert*_(*t*) (units of mV s^*−*1^). The first term *I*_0_ is a constant term representing input to the E or I neuron from other brain areas. For each neuron, *I*_0_ it is drawn from a uniform distribution in the interval [*I*_0*α*_(1*−a*_0_), *I*_0*α*_(1 + *a*_0_)], where *I*_0*α*_ = *N*_*ext*_*J*_*α*0_*r*_*ext*_ (for *α* = *E, I*), *N*_*ext*_ = *n*_*E*_*Np*_*EE*_, and *a*_0_ = 2.5%. *I*_*pert*_(*t*) represents the time-varying source perturbation (see below). The recurrent term evolved according to

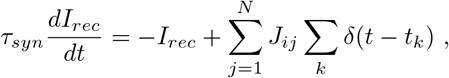

where *τ*_*s*_ is the synaptic time, *J*_*ij*_ are the appropriate recurrent couplings and *t*_*k*_ represents the time of the k-th spike from the j-th presynaptic neuron. Parameter values are in Table III.

**TABLE III.** Parameters for the spiking network in Fig. 5.

| Spiking network simulations |  |  |
| --- | --- | --- |
| Parameter | Description | Value |
| $j_{EE}$ | mean E-to-E synaptic weights $\times \sqrt{N}$ | 0.24 |
| $j_{IE}$ | mean E-to-I synaptic weights $\times \sqrt{N}$ | 0.45 |
| $j_{EI}$ | mean I-to-E synaptic weights $\times \sqrt{N}$ | 3.07 |
| $j_{II}$ | mean I-to-I synaptic weights $\times \sqrt{N}$ | -.15 |
| $J_{E0}$ | mean external input weights to E neurons $\times \sqrt{N}$ | 0.058 |
| $J_{I0}$ | mean external input weights to I neurons $\times \sqrt{N}$ | 0.052 |
| $J_{EE}^+$ | E-to-E within-cluster weight potentiation factor | 31.5 |
| $J_{IE}^+$ | mean E-to-I synaptic weights between clusters | 9.3 |
| $J_{EI}^+$ | mean I-to-E synaptic weights between clusters | 8.6 |
| $J_{II}^+$ | mean I-to-I synaptic weights between clusters | 6.2 |
| $p_{EE}$ | mean E-to-E connection probability within clusters | .2 |
| $p_{IE}$ | mean E-to-I connection probability within clusters | .5 |
| $p_{EI}$ | mean I-to-E connection probability within clusters | .5 |
| $p_{II}$ | mean E-to-E connection probability within clusters | .5 |
| $r_{ext}$ | Average baseline source rate to E and I neurons | 5 spks/s |
| $V_E^{thr}$ | E threshold potential | 1.43 mV |
| $V_I^{thr}$ | I threshold potential | 0.74 mV |
| $V^{reset}$ | E and I reset potential | 0 mV |
| $\tau_m$ | E and I membrane time constant | 20 ms |
| $\tau_{refr}$ | E and I absolute refractory period | 5 ms |
| $\tau_s$ | E and I synaptic time constant | 5 ms |

We modeled the effect of perturbations as a 100ms-long constant step input increase to the external current *I*_*pert*_. Simulations were performed using custom software written in Python. Simulations in the resting periods comprised 8s and in the stimulated protocol to 40s. Each network was initialized with random synaptic weights and simulated with random initial conditions in each trial. Python code to simulate the model during resting periods and perturbations is located at https://github.com/amin-nejat/CCM.

### c. Functional causal flow estimation

The algorithm used for functional causal flow estimation was based on the convergence cross-mapping (CCM) method, proposed in ecosystem analysis [21] and only recently tested on *in vitro* electrophysiology data [26].

In contrast to more traditional causality-detection algorithms based on information transfer, which test the prediction of a downstream time series through information from an upstream one, CCM operates through “nowdiction” (reconstruction of simultaneous time segments) of an upstream series through information from a downstream one. The functional causality relation between any pair of units is then determined by comparing the accuracy of nowdiction in the two directions. As ensured by a powerful theorem [24], nowdiction of the upstream channels tends toward zero error in the asymptotic limit of infinite data size.

Each time segment of data is encoded as a so-called delay vector, by choosing an embedding dimension *d* and a delay time *τ* and constructing higher dimensional vectors

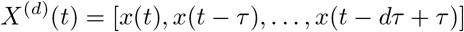

from the given time series *x*(*t*) (Fig. 1). The high-dimensional time series constructed through embedding lives on a manifold diffeomorphic to the full attractor only as long as the embedding dimension *d* is larger than twice the dimensionality of the attractor [23], a statement that generalizes to the box-counting dimension for fractal attractors [24]. This can be far smaller than the number of variables involved in processing, as routinely happens in brain activity during any given task. Two parameters are thus involved in the embedding procedure, the embedding dimension *d* and the delay time *τ*. The condition that *d* be large enough is sufficient for an ideal setting, but how to select *d* and *τ* for a given real dataset has been the topic of a vast literature (see [35]). The choice of *τ* and *d* depend on the specific dataset. For the spiking data from alert monkeys and network simulations, we estimated spike counts in *b* = 60*ms* bins, used a delay time *τ* = *b* and an embedding dimension *d* = 20, although we confirmed that reconstruction results held robustly for a wide range of *d* (not shown). We split the full multi-dimensional time series into two segments – a training period and a test period. Specifically we choose the latter from the end of the sample being studied, and took it to be 1/10 of the full. Given that cross-validation serves as a guarantee against overfitting, it allows us to rely on a simple nearest neighbor algorithm to concretely perform reconstructions.

Given two time series *x*(*t*) and *y*(*t*), d-dimensional time series of delay vectors *X*(*t*) and *Y* (*t*) are constructed. To test the accuracy of reconstruction, the data are segmented into a library period and a test period. For each putative downstream vector *X*(*t*) in the test sample, a reconstruction *Ŷ* (*t*) is obtained by listing the *k* time points *t*_*j*_[*t*] (*j* = 1, …, *k*) corresponding to the delay vectors that are nearest neighbor to *X*(*t*) according to the euclidean distance Δ_*j*_(*t*) =||*X*(*t*) *X*(*t*_*j*_[*t*])||. For each neighbor, the corresponding weight is computed as a positive, decreasing function of its distance from *X*(*t*), namely *w*_*j*_(*t*) = *f*_*j*_(Δ_1_(*t*), …, Δ_*k*_(*t*)), normalized to yield the reconstruction

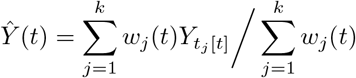

In the limit of infinitely long datasets, any finite *k* and any function *f* will yield asymptotically the same reconstruction. For a finite dataset, a common choice which we adopted is a uniform weight associated to the “simplex dimension” *k* = *d* + 1 [112]. The major computational bottleneck lies in the extraction of the nearest neighbors.

We used a ball tree data structure, which partitions data in a series of nesting hyper-spheres as suitable to the structure of the training data. Reconstruction for a single recording took a matter of minutes on a regular laptop.

Once reconstructions are obtained, their accuracy is estimated through the Pearson correlation coefficient *r* between the test-period time series *Y* (*t*) and its reconstruction *Ŷ* (*t*). In the noiseless infinite-data limit such correlation saturates to one if a causal flow exists in the direction opposite to reconstruction. With finite sample size, we adopted significance of the correlation coefficient as a sufficient condition for various causal scenarios – unidirectional flow in either direction, recurrence, and independence (Fig 2). Notice that the chosen measurement of reconstruction in terms of a Pearson correlation *r* is a random variable over the space of system trajectories but not necessarily a normal one – being a correlation coefficient it is in fact bounded between −1 and 1. This poses statistical problems to comparing different coefficients, and in particular coefficients in two converse directions ([113, 114]). Crucially, for this reason we define the FCF as the Fisher z-transform of the reconstruction coefficient *r*, which can be relied upon to have near-normal distribution [34].

#### 1. Data preprocessing

While the creation of delay vectors directly from spikes has been explored in the literature [115], we found it convenient to use spike counts as continuous variables from whose time series we built delay vectors. Smoothing the spike counts is a delicate step that can introduce extraneous interference even if the linear filter is of a causal type. Specific denoising techniques [116] have been proposed to circumvent this problem for the preprocessing pipeline of delay-embedding analyses. We found that the least invasive approach was to rely entirely on a nearest neighbor method to perform the denoising.

Since the binning already performs a temporal coarse graining on the information available from recordings, we chose to pick the delay time step equal to one in units of the bin width *b* = 60ms. The bin width and all the other hyperparameters are listed in table IV.

**TABLE IV.** Hyperparameters applied to estimate causal flow from resting period spiking activity in awake monkeys and network simulations.

| Hyperparameters for spiking activity |  |  |
| --- | --- | --- |
| Parameter | Description | Value |
| $b$ | bin size for spike counting | 60 ms |
| $\alpha$ | ratio of recording used as test sample | 0.1 |
| $\tau$ | delay time (in units of $b$) | 1 |
| $d$ | embedding dimension | 20 |
| $k$ | number of nearest neighbors for reconstruction | 20 |

Properties studied by delay embedding are invariant under any differentiable change of variables, and in particular under linear transformations. However, because differences in scale between the activities of individual neurons can affect the calculation of nearest-neighbor weights, the firing rate time series of all channels were shifted and normalized to have zero mean and unit variance, thereby removing concerns about the scale dependence of the exponent in the nearest neighbor weights.

### D. Significance of causal flow

To establish significance of estimated values of FCF, we adopted an approach to hypothesis-testing now widely used in nonlinear science that consists in generating “surrogate” data, i.e. artificially constructed time series that match the original dataset according to some statistical benchmarks but where the property being tested has been scrambled. The ranking of a discriminating statistic over the distribution of the same quantity calculated on the surrogate allows a significance test on the hypothesis. We chose a surrogate generation method, first proposed in [35], that was designed to preserve all largescale nonlinear properties of the system. Surrogate time series are produced in three stages. Firstly, we evaluated phase-space distances among Takens states constructed from each time series: nearest neighbors were defined as states within a certain maximum radius from each others. Secondly, an equivalence relationships is defined between states possessing the same set of neighbors (known as “twins”). Finally, surrogate trajectories are initialized randomly and generated by allowing each subsequent step to start with equal probability from the state it just reached or from one of its twins. Each set of twins is thus replaced by a probability superposition of them – macrostates that coarse-grain phase space by making trajectories stochastic. Each surrogate time series emerges thus as the instantiation of a Markov process whose transition matrix has diagonal elements *p*_*ii*_ = (*n*_*i*_*−*1)*/n*_*i*_, for *n*_*i*_ equal to the number of twins. This method has the advantage of preserving the temporal statistics of the full system, at the price of introducing a hyperparameter, the neighborhood radius, choosen as the 10% quantile in the nearest-neighbor distances distributions of Takens states as the threshold for neighborhood to base the twinning upon.

The rationale for choosing the twin surrogate method described above as opposed to surrogates based on random reshuffling of time points (which destroys all but the amplitude distribution) or isospectral surrogates (based on reshuffling the phases of the Fourier transform and anti-transforming with certain precautions about boundaries, thus preserving autocorrelation) is the following. The latter two methods destroy not just the causal links but rather any nonlinear property of the system (e.g. its density distribution in phase space and its entropy) and therefore pose a considerable risk for false positive. We thus adopted the choice of twin surrogates as a vastly more conservative one which preserves the nonlinear properties of the system while destroying the causal links (for details see [35]).

#### 1. Estimating perturbation effects

We estimated perturbation effects by comparing network activity in intervals of length *T*_*max*_ ending Δ_*pre*_ before the onset and beginning Δ_*post*_ after the offset of each perturbation. Such cushion periods reflected the transient pause in recording in those short periods adjacent to each microstimulation (see below). The spiking activity of each target neural cluster was estimated in *dT* bins (see Table V) covering the lapse interval *T*_*max*_ and the distribution of spike counts was aggregated across all stimulation trials of the same source. The results were largely consistent for a variety of bin sizes and *T*_*max*_; we chose values that maximized our analysis power without obscuring the dynamics of the effects. A Kolmogorov-Smirnov (KS) test between the pre- and post-stimulation aggregated spike count distributions was performed to assess a significant effect of a perturbation, whose effects were reported in the form of KS statistics and p-value (Fig. 6 and Fig. 7). All parameters are reported in Table V.

**TABLE V.**
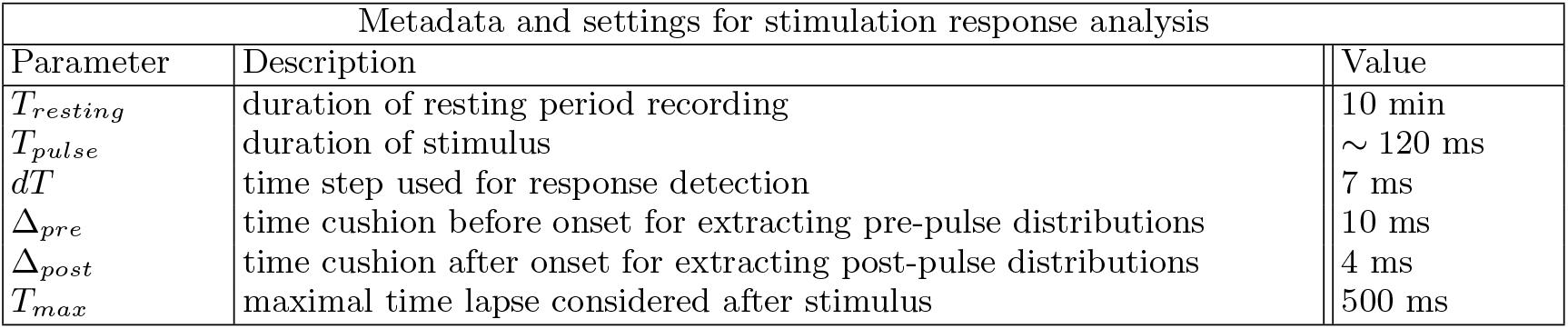
Metadata and settings for response analysis.

### E. Experimental data

We recorded and perturbed the activity of neurons in the pre-arcuate gyrus (area 8Ar) of macaque monkeys (macaca mulatta) using chronically implanted multielectrode Utah arrays (96 electrodes; Blackrock Microsystems). All experimental procedures conformed to the National Institutes of Health *Guide for the Care and Use of Laboratory Animals* and were approved by the New York University Animal Welfare Committee.

During the experiments, monkeys sat in a primate chair, with their heads fixed using a titanium head post. The room was mildly lit and quiet. Monkeys did not perform any task nor receive reward. We monitored the monkey’s eye and limb movements using infra-red camera systems (Eyelink for eye tracking, 1 KHz sampling rate). The monkey remained awake (open eyes) and rarely moved limbs during these resting blocks, which were typically 10-20 min long. We started a session with a resting period recording block that we used to quantify functional causal flows and predict the perturbation effects. This block was followed with a recording and microstimulation block that we used for testing the predictions.

The electrodes of the Utah array were 1mm long with 400 *μm* spacing between adjacent electrodes, permitting simultaneous recordings from neighboring columns in a 4 mm*×*4 mm region of cortex. Raw voltage signals were filtered and thresholded in real time to identify spikes. Spike waveforms and raw voltage were saved at 30 KHz sampling frequency for offline processing.

Electrical microstimulation was delivered through individual electrodes of the Utah array. Microstimulation pulse trains consisted of low current (15 *μA*) biphasic pulses [2, 3, 92], each 0.2 ms long, delivered at 200 Hz. Pulse trains were 120 ms long and occurred once in any 5 s period. The exact time of the microstimulation with the 5s periods varied randomly. Electrophysiological recording was done in between the microstimulation trains; it resumed with a short latency (¡5 ms) at the end of each pulse train and continued until the beginning of the subsequent train. Microstimulation of the prearcuate gyrus with currents ¿50 *μA* could trigger saccadic eye movements [117]. Our low current microstimulation was chosen well below this motor threshold and never triggered saccades in our experiments.

### F. Spatial dependence of FCF and perturbation

Both the FCF and the perturbation effects decayed with the distance from the source electrode. In order to control for this effect, we performed a partial correlation analysis by detrending the spatial dependence of FCF and the perturbation effects (KS) using a linear regression, and then reevaluating the Pearson correlation between the their residuals (Fig. S4). In particular, fixing a source *j* and given three measurements for each target *i*, namely *FCF*_*i*_ (functional causal flow), *KS*_*i*_ (interventional connectivity, and *d*_*i*_ (physical distance between electrodes *i* and *j*) we fit two linear regression models:

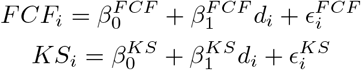

The partial correlation between FCF and KS controlling for physical distance is then given by: *ρ*(*ϵ*^*F CF*^, *ϵ*^*KS*^). This quantity allows us to measure the unique contribution of FCF in predicting KS by removing their linear spatial dependence. We further repeated this experiment nonparametrically by subtracting the median curve shown in Fig. 7 and computing the correlation between the residuals. These results confirm our observation that the relationship between FCF and KS is not simply a confound of the physical distance between the electrodes.

### G. Comparison to Granger Causality

We adopted the likelihood ratio as our Granger statistic.

For the conceptual comparison in panel A of Fig. S5, we considered the straightforward solution to the linear equations

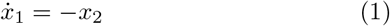

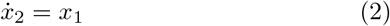

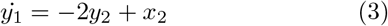

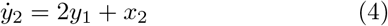

and added white zero-mean Gaussian measurement noise of standard deviation *σ* = .1 on top of all four variables. Causality between *x*_2_ and *y*_2_ was inferred with FCF and Granger Causality, using the same significance criterion (*p <* 0.05) for both directions and for both methods. Moreover the same value *d* = 10 was used for both the maximal lag in Granger and the delay dimension for FCF (adopting, accordingly, delay time *τ* = 1).

In Fig. S5, panel B, the variable representing single-cell activity was taken to be the x-coordinate of a Lorenz attractor with parameters *α* = 10, *β* = 8*/*3, *ρ* = 28, while the activity of the complex cell donwstream of it evolved as 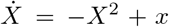. Causality was inferred with corresponding parameters in the two method, exactly as done for panel A.

The Granger analyses of pseudodata from simulations presented in Fig. S6, as well as the analyses on alert monkey data presented in Fig. S7, were implemented using custom-made Python code based on the Matlab package of ([118]). The package can in principle implement both standard GC which operates separately on individual pairs just as FCF, and multivariate GC (“MVGC”) which conditions away influence from thirdparty units. It is worth noting that, depending on the dataset, the conditioning operated by MVGC could improve or worsen the ability to single out causal links. It can worsen performance when units share commons causal input travelling at different speeds to the system; if the causal connection from A to B is slow and the one from A to C is fast, discounting the effects of C can lead to the erroneous conclusion that A is not causing B.

## SUPPLEMENTARY MATERIALS

**FIG. S1.**
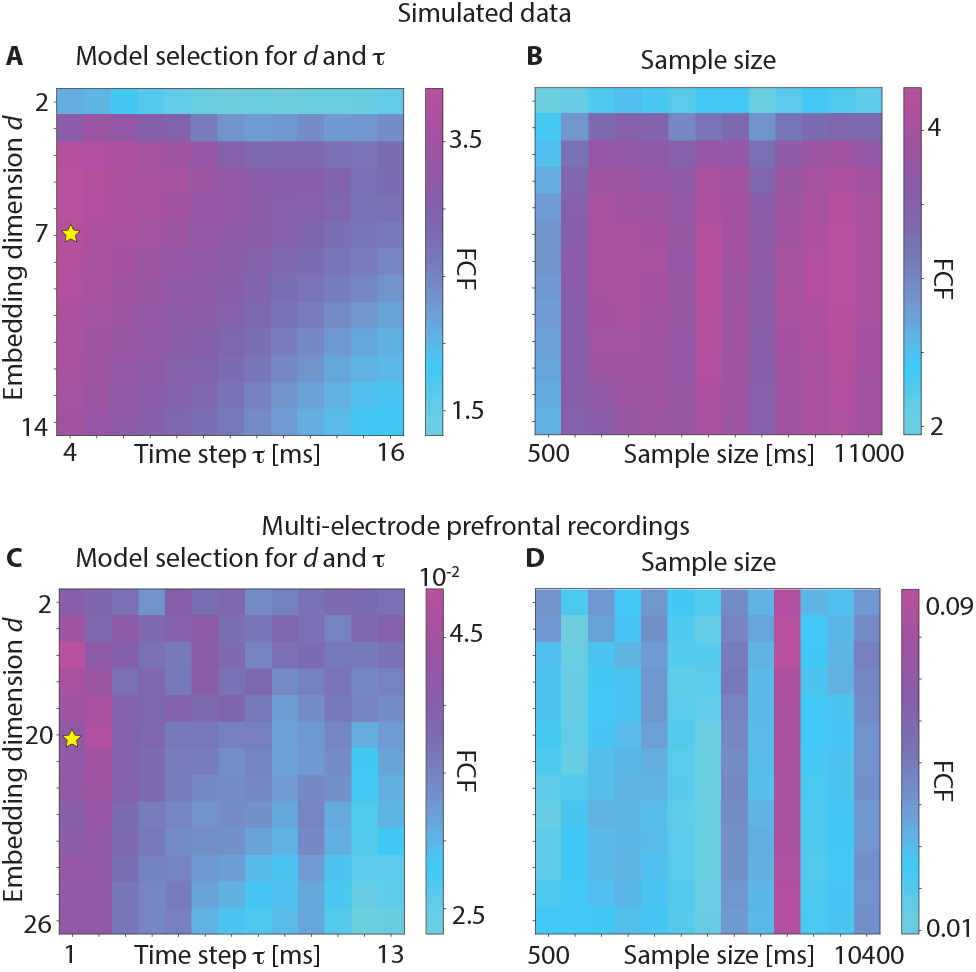
Model selection and sample size dependence. The FCF hyperparameters *d* (embedding dimension) and *τ* (time step) determining the delay vectors [*x*(*t*), *x*(*t − τ*), …, *x*(*t − dτ* + *τ*)] used for reconstruction were chosen by a hyperparameter search in the simulated continuous rate network (A) and in the electrophysiological data from alert monkeys (C). The interaction between embedding dimension *d* and sample size used for FCF inference in the simulated continuous rate network (B) and in the electrophysiological data from alert monkeys (D).

**FIG. S2.**
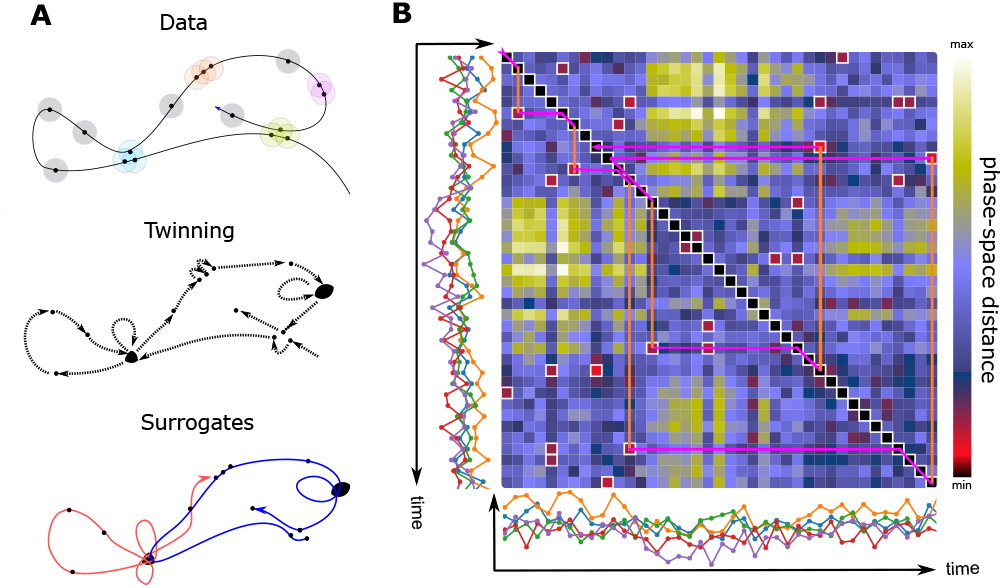
Establishing significance of functional causal flow (FCF). A) Significance of FCF is determined by comparison to surrogate datasets designed to preserve all large-scale non-linear properties of the system. Surrogates are produced in three stages: top, phase-space distance is evaluated among Takens states constructed from each time series, and nearest neighbors are identified; center, states in the trajectory are coarse-grained by collapsing states with the same set of neighbors (in the example, the blue and purple clusters merge but the orange and green ones do not); bottom, surrogate trajectories are generated from random initial conditions by regarding twin-sets as retentive states in a Markov process (retention is represented by self-loop and has probability *p* = (*n −* 1)*/n* where *n* is the numebr of twins). B) Example trajectory depicted over the matrix of phase-space distances for the multi dimensional time series shown along both axes. Whereas the main diagonal corresponds to the flow of the recorded time series, at each step the surrogate time series can either move forward as in the recording or depart from the main diagonal (vertical orange lines) to pick one of the low-distance states in its own twin set, and perform from there a forward step mimicking the recording (purple broken lines). Notice that, as in the example, motion can be both forward and backward in time.

**FIG. S3.**
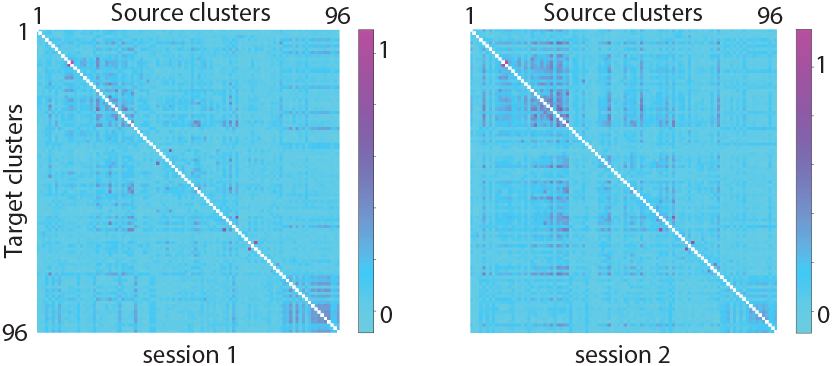
FCF between electrodes in the monkey prefrontal cortex. Full resting state FCF matrix was inferred from ensemble spiking activity recorded by multi-electrode arrays in the pre-arcuate gyrus during quiet wakefulness. Two sessions are shown (same sessions as in Fig. 7).

### A. Comparison to Other Causality Indices: Conceptual differences

In the main text, we compared the performance of FCF in predicting perturbation effects with that of several alternative methods based on information theory, including: Granger causality in its univariate (GC) and multivariate versions (MGC); nonlinear GC (NGC) and extended GC (EGC), which perform autoregression using radial basis functions; and transfer entropy (TE). These methods rely on an assumption of “separability” between cause and effect, i.e., information on the upstream variable being transferred but not stored into the downstream variables. There are clear indications of this assumption breaking down in neurophysiological recordings [119]. This breakdown can occur even in the simplest systems. In panel A of Fig. S5, the 2D trajectory of an upstream variable (say, a network’s activity) moves along a linear cycle with measurement noise on top. A downstream variable linearly integrates input from this upstream variable and is overlaid with extra measurement noise (see Methods for details). Application of GC – which assumes separability – detects causation in both directions, including the wrong one. By contrast, FCF infers causality only in the correct direction.

**FIG. S4.**
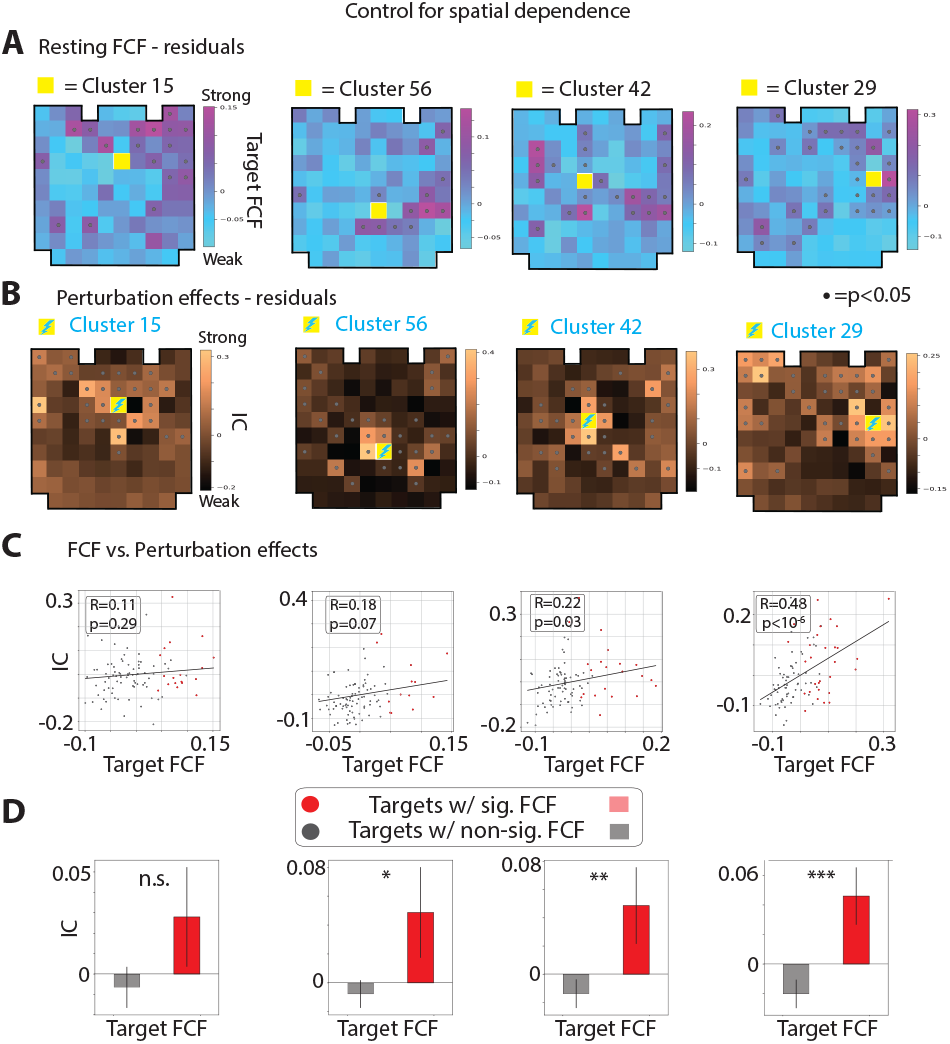
Controlling for spatial dependence of FCF and perturbation effects. A-B) The residual FCF (A) and perturbation effects (B; quantified with interventional connectivity) from Fig. 7A-B. Residuals are calculated by removing the dependence on distance from the source electrode. C) Correlation between FCF and interventional connectivity after controlling for spatial dependence. D) Mean residual IC for target clusters with significant and non-significant FCF for each source electrode. Conventions are similar to Fig. 7.

The nested loop inside the trajectory of the downstream variable is a quintessential example of “downstream complexity” [27]. Because of that extra complexity, the mapping inference that FCF attempts is less accurate in the downstream direction. GC, on the contrary, is concerned with detecting signal over noise and, regardless of the direction, is able to receive help from information in the source variable. Knowing the recent past of the downstream trajectory helps predicting the next step of the upstream trajectory, which aside from the loop is equivalent.

**FIG. S5.**
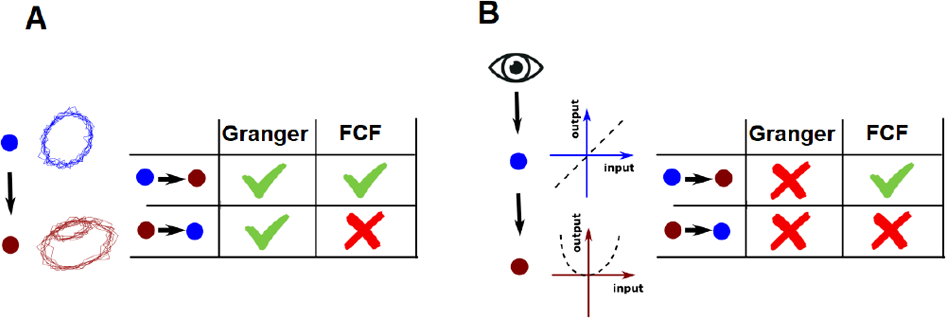
Conceptual comparison of Granger Causality to Functional Causal Flow. Panel A) In the presence of nonseparable dynamics, application of Granger causality detects causation in both directions, including the wrong one. By contrast, FCF infers causality only in the correct direction. Panel B) Nonlinear interaction as represented by a complex cell quadratically integrating signal from a simple cell. Granger causality here detects no causality in either direction. By contrast, FCF correctly infers causality only in the downstream direction.

GC also assumes monotonicity in causation trends, which cannot be expected to occur with real neurons. Consider (Fig. S5, panel B) the typical case of external stimulus integrated linearly by a simple cell whose output is integrated by the nonlinear input-output function of a downstream complex cell (Methods for details). This basic architecture is already beyond the range of validity of Granger causality, which here detects no causality in either direction. By contrast, FCF correctly infers causality only in the downstream direction.

### B. Comparison to Other Causality Indices: Results

A recent paper [120] investigates how different causality indices recover the direction of causation in simulations where there is a clear unidirectional influence from one variable onto the other. Here we apply those indices to our simulations and real data to address whether or not other indices can recover the direction of influence in the presence of recurrence and network dynamics. Below we first briefly explain each causality index. Then, we present results on the simulated rate network from Figs. 2-4 and monkey prefrontal data from Fig. 7 to investigate which indices can predict perturbation effects measured by interventional connectivity.

A general principle underlying all alternative causality indices is that causality is defined by the precedence of influence in time. If the past of variable *X* contains information about or allows the prediction of the future of variable *Y* then there is causal influence from *X* to *Y*. This is precisely the idea behind the definition of *Granger Causality (GC)* and its variants. If we assume that two signals evolve jointly according to an autoregressive model, then GC measures if the past of *X, Y* together helps predicting the future of *Y* better than the past of *Y* alone. The significance test is performed using F-test as commonly done in the GC literature.

*Transfer Entropy (TE)* is defined similarly, but TE relaxes the autoregressive assumption to arbitrary rules for the stochastic evolution of time series, computing the conditional mutual information between the future of *Y* and past of *X* conditioned on the past of *Y* [121, 122]. It is worth noting that if the data follows an autoregressive model, GC and TE become equivalent. Although TE is nonparametric, its estimation is a challenging statistical task often requiring large amounts of data. For TE, here we use an estimator developed by [123] and employed by [120] which is based on nearest neighbor methods.

Although GC is originally developed for univariate and autoregressive signals, one can generalize it to multivariate and nonlinear counterparts. *Multivariate GC* (MGC) computes the same criterion as GC with the difference that the conditioning is done on all the other variables in the multivariate time series. This allows for measuring the unique predictability of future of *Y* from past of *X* when we control for other intermediate signals in the network. *Nonlinear GC* (NGC) performs the autoregression using radial basis functions [124]. *Extended GC* (EGC) provides another generalization to GC based on locally linear approximation [125].

Notice that in contrast to these methods which rely on the stochastic fluctuations of signals, FCF is based on the deterministic aspects of a dynamical system and instead of measuring noise statistics it uses nearest neighbors in the state space of a stationary dynamical system to predict one signal from the other. Moreover, in the limit of large datasets, FCF is less affected by the unobserved nodes due to the topological correspondence between the time-lagged history of each variable and the high-dimensional data generating system which includes the unobserved nodes.

We summarize the results obtained from different causality indices in Figs. S6 and S7. In the simulated rate network, FCF and GC show significant positive correlations with IC, and FCF performs best. For the monkey data, GC and FCF show positive correlations with IC but FCF is more robust and outperforms other indices. TE fails perhaps due to the small sample size or the presence of observed and unobserved nodes in the network which are not accounted for. A summary of the hyperparameters used for the simulations and calculating different causality indices are included in the corresponding `config` file on the code repository released with this paper: https://github.com/amin-nejat/CCM/tree/master/example_configs.

**FIG. S6.**
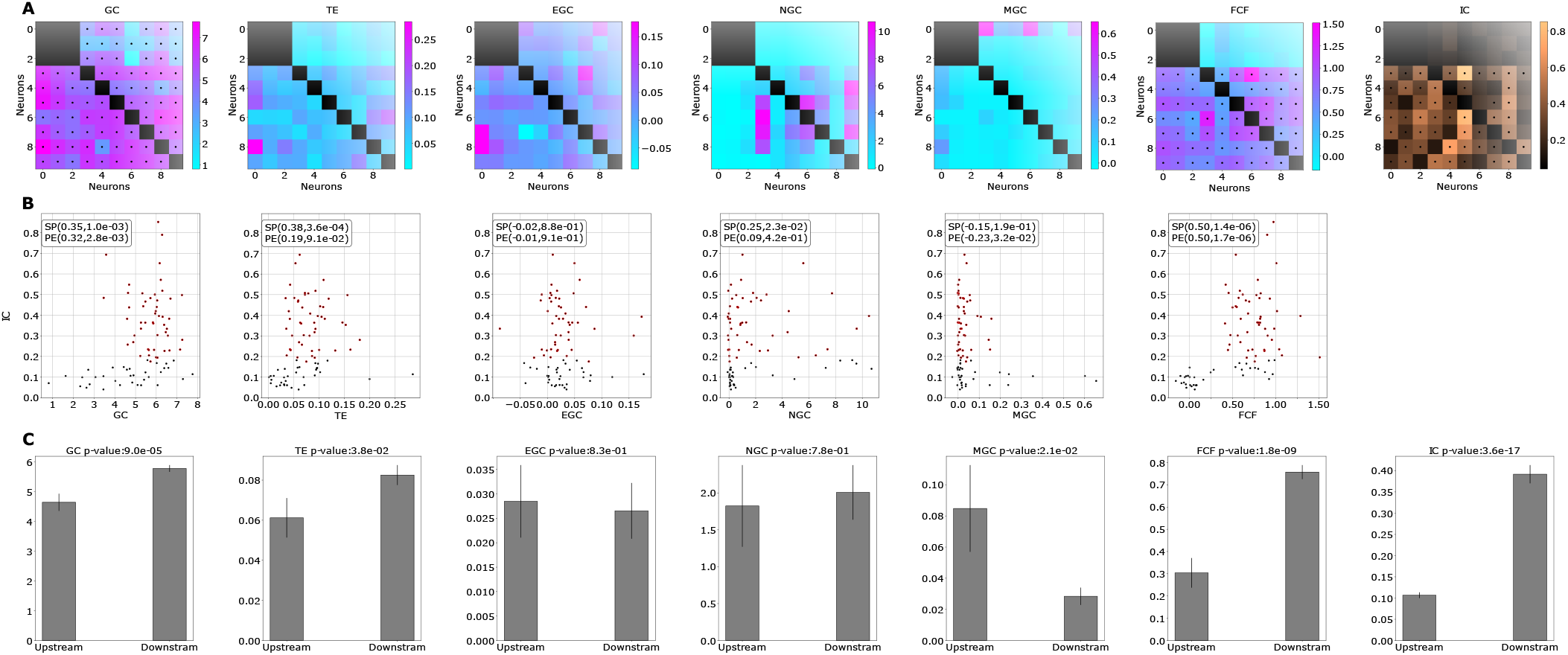
Comparison of causality indices on the simulated rate network: A) Different causality indices applied to the simulated rate network of Fig. 2-4; from left: Granger Causality (GC), Transfer Entropy (TE), Extended Granger Causality (EGC), Nonlinear Granger Causality (NGC), Multivariate Granger Causality (MGC), Functional Causal Flow (FCF), Interventional Connectivity (IC). B) Scatter plots of correlations between measured causality index on the x-axis and interventional connectivity values on the y-axis (SP, PE represent Spearman and Pearson correlation coefficients and p-values, respectively). In this simulation FCF can best predict IC among the indices. C) Each bar plot corresponds to the causality index values separated according to the significance of IC matrix (t-test with respective p-values reported); FCF best reflects upstream vs. downstream as defined by the significant elements of IC. The rightmost bar plot corresponds to the IC values separated by their significance, providing a ceiling for the causality indices.

**FIG. S7.**
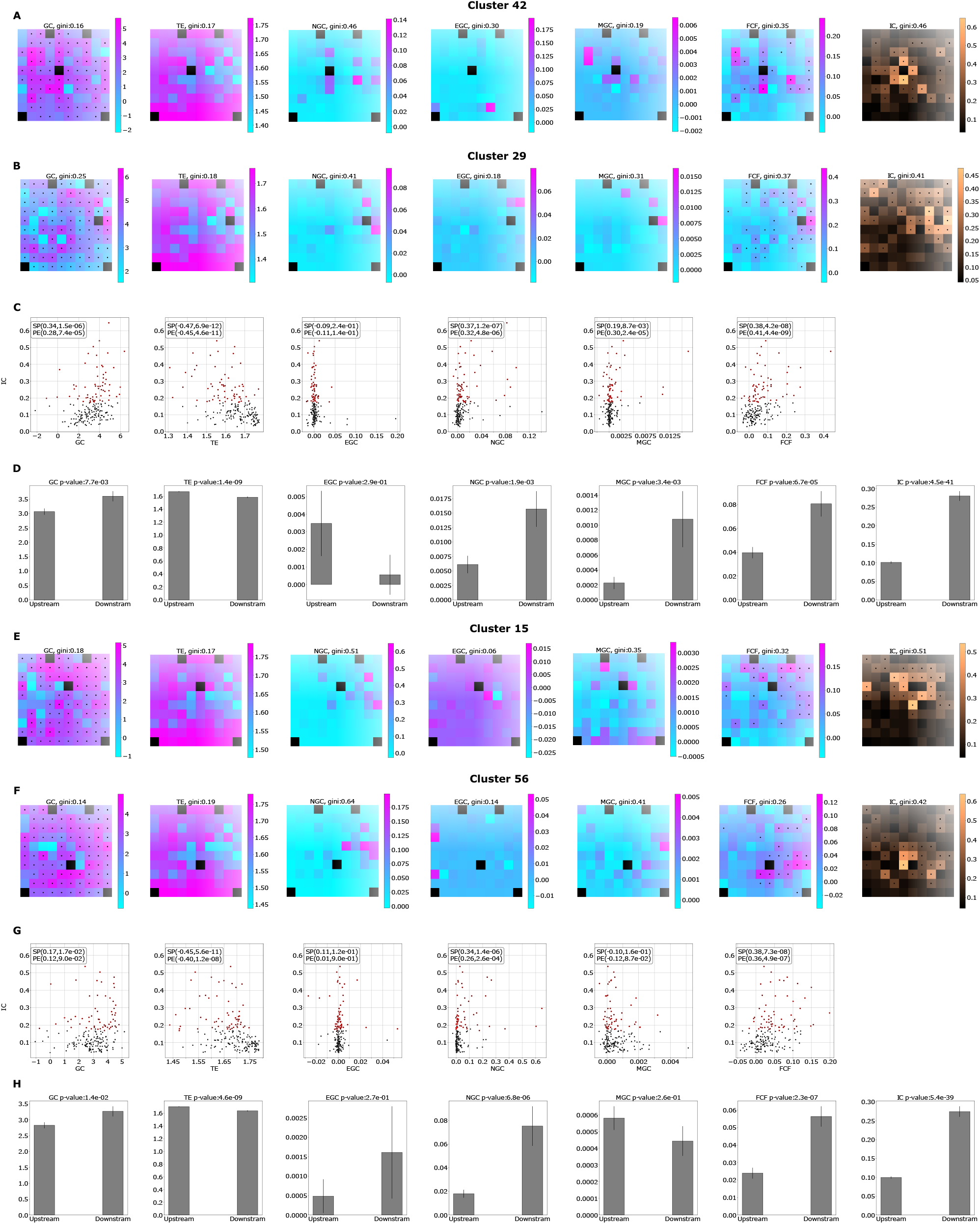
Comparison of causality indices on the monkey data: A, B, E, F) Causality indices computed on two stimulated channels during the resting activity (see Fig. S6 for notations); each image corresponds to the causality between the stimulated channel and all other channels organized in the physical layout of the electrode array (same as in Fig. 7; Gini index of causal vector reported on top). C, G) Scatter plots of correlations between measured causality index on the x-axis and interventional connectivity values on the y-axis (SP, PE represent Spearman and Pearson correlation coefficients and p-values, respectively), in this dataset FCF can best predict IC among the indices. D, H) Each bar plot corresponds to the causality index values separated according to the significance of IC matrix; FCF best reflects upstream vs. downstream as defined by the significant elements of IC. A-D and E-H correspond to the two different recording sessions, respectively.

